# The role of SOX9 in non-small cell lung cancer progression is histopathology-selective

**DOI:** 10.1101/2020.11.23.393926

**Authors:** Jie Bao, Katja Närhi, Ana Teodòsio, Annabrita Hemmes, Nora M Linnavirta, Mikko I Mäyränpää, Kaisa Salmenkivi, John Le Quesne, Emmy W Verschuren

**Affiliations:** Institute for Molecular Medicine Finland, HiLIFE, University of Helsinki, Helsinki, Finland; GlaxoSmithKline, Espoo, Finland; MRC Toxicology Unit, University of Cambridge, United Kingdom; HUSLAB, Division of Pathology, Helsinki University Hospital, Helsinki, Finland; Department of Pathology, University of Helsinki, Helsinki, Finland; Leicester Cancer Research Centre, University of Leicester, Leicester, United Kingdom

**Keywords:** SOX9, non-small cell lung cancer, histopathology, cell of origin, metastasis, extracellular matrix, collagen IV, *in vivo* models

## Abstract

The transcription factor SOX9 is a key regulator of multiple developmental processes, and is frequently re-expressed in non-small cell lung cancer (NSCLC). Its precise role in the progression of NSCLC histopathologies has however remained elusive. We show that SOX9 expression relates to poor outcome and invasive histopathology in human adenocarcinomas, and is absent in murine early minimally invasive and human *in situ* adenocarcinoma. Interestingly, despite wide SOX9 expression across advanced NSCLC histotypes, its genetic deletion in the murine *Kras^G12D^;Lkb1^-/-^* model selectively disrupted only the growth of papillary NSCLC, without affecting the initiation of precursor lesions or growth of mucinous or squamous tissue. Spatial tissue phenotyping indicated a requirement of SOX9 expression for the progression of surfactant protein C-expressing progenitor cells, which gave rise to papillary tumours. Intriguingly, while SOX9 expression was dispensable for squamous tissue formation, its loss in fact led to enhanced squamous tumour metastasis, which was associated with altered collagen IV deposition in the basement membrane. Our work therefore demonstrates histopathology-selective roles for SOX9 in NSCLC progression, namely a requirement for papillary adenocarcinoma progression, but opposing metastasis-suppressing function in squamous histotype tissue. This attests to a pleiotropic SOX9 function, linked to the cell of origin and microenvironmental tissue contexts.

## Introduction

Lung cancer remains the leading cause of cancer-related mortality, the majority of which represent non-small cell lung cancer (NSCLC; 85%). NSCLC is subdivided via its two major histotypes, adenocarcinoma (AC) and squamous cell carcinoma (SCC), of which the AC group is most heterogeneous [1–4]. ACs are further subdivided based on the tissue’s predominant histotype structure, particularly papillary AC (PAC), mucinous AC (MAC), and acinar AC (AAC) [5]. Certain driver mutations are selectively enriched in these AC subtypes. For example, *KEAP1* mutation or *NRF2* overexpression occurs more frequently in PACs when compared to total NSCLCs, and *KRAS, EGFR* or *ALK* mutations have respectively been associated with mucinous, lepidic growth, or signet ring structure [6, 7]. Molecular and genetic functions therefore need to be interpreted within these histopathology-specific tissue contexts.

The transcription factor SOX9 is a member of the SOX (sex-determining region Y-related) family [8], and regulates a variety of developmental processes [9–11]. During murine lung development, SOX9 is expressed in a timed manner to coordinate lung branching morphogenesis and alveolar differentiation [12, 13]. SOX9 is silenced in adult tissues but frequently re-expressed in cancer, where it has been associated with increased cell proliferation, transcriptional reprogramming and self-renewal of stem cells, and increased cell invasiveness and drug resistance [14–17]. The *SOX9* gene resides on the long arm of chromosome 17 (17q24.3), which is frequently gained in non-smokers with lung AC [18]. In patients, elevated SOX9 expression has been associated with poor survival and advanced clinical NSCLC stage, and its expression was shown to promote the proliferation of AC cells *in vitro* through affecting the CDKN1A and CDK4 cell cycle regulators [19, 20]. However, while many studies find oncogenic roles for SOX9, supporting its therapeutic targeting, also the downregulation of SOX9 has been linked to relapse of prostate cancer [21] and stage II colon cancer [22], indicating that SOX9 has pleiotropic functions. Despite NSCLC constituting a heterogeneous set of diseases, the potential histopathology-selective role of SOX9 in NSCLC progression has thus far not been studied.

Distinct sets of developmental signals and transcription factors are expressed in different lung progenitor cells, and thereby exert lineage-specific functions [23]. In recent years, the ways in which different cells of origin influence NSCLC progression have been addressed by applying genetically engineered mouse models [24]. Using adenovirus-mediated Cre-recombination, we previously showed that conditional *Kras^G12D^* expression combined with loss of *Lkb1* in bronchial progenitors expressing Clara cell antigen 10 (CC10) gives rise to a wide NSCLC histotype spectrum encompassing minimally invasive AC (IAC), MAC, AAC, PAC as well as adenosquamous carcinoma (ASC) [25], a rare NSCLC subtype exhibiting both SCC and AC features [26]. To the contrary, in alveolar type II (AT2) progenitors expressing surfactant protein C (SPC), the same genetic alterations mainly lead to IAC and PAC formation [25]. To study the histopathology-selective roles of SOX9 expression in NSCLC, we established heterozygous and homozygous conditional *Sox9* null *Kras^G12D^;Lkb1^-/-^* cohorts, and examined how *Sox9* loss influenced the progression of lesions of distinct histotypes. We describe opposing histopathology-selective roles of SOX9 in NSCLC progression and metastasis, and link these to the cell of origin and tumour microenvironment contexts, with ramifications for future directions in therapeutic intervention.

## Materials and Methods

### Human and murine tumour tissues

Breeding of *Kras^G12D/+^;Lkb1^flox/flox^ (Kras^G12D^;Lkb1^-/-^*) mice, genotyping, and lung tumour initiation by intranasal infection with progenitor cell-directed Ad5-Cre viruses were performed as previously described [23]. *Kras^G12D^;Lkb1^-/-^;Sox9^+/-^* and *Kras^G12D^;Lkb1^-/-^;Sox9^-/-^* cohorts were generated by crossing *Kras^G12D^;Lkb1^-/-^* and *Sox9^flox^* mice (*B6.129S7-Sox9^tm2Crm^/J*, The Jackson Laboratory). *Sox9* genotypes were determined using the supplied protocol 29713 (standard PCR Assay - Sox9<tm1Crm>, The Jackson Laboratory). Animal handlings and studies were performed according to the guidelines from the Finnish National Board of Animal Experimentation, approved by the Experimental Animal Committee of the University of Helsinki and the State Provincial Office of Southern Finland (ESAVI/6365/2019). Formalin fixed paraffin embedded (FFPE) samples of human NSCLC SCC, ASC and PAC lesions were collected from the Helsinki Biobank’s pathology sample archive (Helsinki Biobank project number HBP2016002), and a tissue microarray (TMA-1) was constructed as previously described [27]. Tumour stages were determined by a certified thoracic pathologist, following American Joint Committee on Cancer (AJCC) criteria. Detailed information and raw data from the murine and human NSCLC sample analyses are provided in the supplementary material, Table S2 and S3. The large human adenocarcinoma TMA collection (TMA-2; 970 samples) was obtained from a continuous retrospective cohort of archival human lung adenocarcinoma tissue from University Hospital Leicester, under research ethics committee approval. Clinicopathological and follow-up data was obtained from regional and national databases, and all morphological assessments were performed by a subspecialty consultant diagnostic histopathologist.

### Tissue processing, immunohistochemistry (IHC) and image acquisition

Murine lung tumours were processed and embedded in paraffin as previously described [23]. FFPE tissue blocks containing whole murine lung or TMA blocks were cut as 4 μm thick sections. Two sequential sections were placed on one glass slide (SUPERFROST® PLUS, Thermo scientific, Ref J1800AMNZ), and labelled to mark the sectioning order. Adjacent tissue sections were each stained with one antibody. Standard IHC was performed with the primary antibodies listed in supplementary material, Table S1. For routine signal development, BrightVision poly-HRP goat anti-rabbit IgG (Immunologic B.V., no dilution) was incubated at room temperature for 30 min. For developing the SOX2 signal, DaG-HRP (Santa Cruz Biotechnology sc2020, 1:100) was incubated at room temperature for 30 min. For signal detection, Bright-DAB Substrate Kit (Immunologic B.V.) was used. Whole slide bright-field scans of stained murine and clinical tissue sections were acquired by a digital slide scanner (Pannoramic 250, 3DHISTECH Ltd) using a 20x objective. 3 μm sections from the TMA-2 human adenocarcinoma cohort were duplex stained using a Ventana autostainer with antibodies for cytokeratin AE1/AE3 (1:250, #NCL-L-AE1/AE3, Leica detected with Roche discovery yellow chromogen) and SOX9 (1:10000, AP5535, Millipore detected with Roche discovery purple chromogen). Up to three 1 mm cores per tumour were analysed. Images of haematoxylin-counterstained images were acquired using a Hamamatsu nanozoomer.

### Region-of-interest (ROI)-specific image analysis

Histopathology classification was performed in conjunction with an expert pathologist as previously described [23, 24]. The sizes of individual tumours and lung tissues were measured using Pannoramic CaseViewer (3DHISTECH Ltd). For image analysis, TIFF images of murine lungs with hematoxylin and eosin (H&E) and biomarker stainings on whole slide scans were exported using the Pannoramic CaseViewer (3DHISTECH Ltd) at a magnification of 1:2. Individual tumours on H&E-stained images were then outlined and annotated by their histotypes using ImageJ/Fiji [28] (‘Analyze’-> ‘Tools’-> ‘ROI manager’) to generate a ROI library for each murine sample. Images of serial sections stained with SOX9, Ki67, pAKT and pERK were registered to the nearby H&E images using our previously reported open-source software Spa-R [29], which allows ROI libraries to be applied to marker-stained images (with slight adjustments). For quantifying Ki-67 and SOX9-positive nuclei, individual tumours were exported as inverted 8-bit grayscale images and processed with a CellProfiler pipeline (www.cellprofiler.org) [30] built with nuclei segmentation, intensity measurement, threshold-based filtration, and percentage calculation modules. The parameters in the pipeline were specified for each staining. To measure pERK and pAKT staining in individual tumours, a workflow using Spa-R and ImageJ/Fiji, and thresholds decided by three independent immunohistochemistry experts, were implemented as before [29]. All raw results can be found in the supplementary material, Table S3. To mark Collagen IV fibres with red pseudo-colour, our in-house open-source Spa-Q software [29] was applied, using identical thresholds for all samples.

### Image analysis of human adenocarcinoma cohort (TMA-2)

SOX9 duplex core images were quantified with a purpose-made Visiopharm app. In brief, images were segmented into epithelial and stromal elements using the yellow channel with manual assistance where necessary. Tumour cell nuclei were categorised as weak, negative, moderate or strong for purple SOX9 staining using human-defined cut-offs, and an H-score was calculated for each core (%weak + 2x%moderate + 3x%strong nuclei, yielding a score from 0-300). Ki-67 staining was performed with the Ventana pre-diluted ready-to-use kit, and assessed by a trained histopathologist.

### Statistical analysis

For clinical sample (TMA-1) and murine sample analysis, data visualisation and statistical analyses were performed with GraphPad Prism 8 (GraphPad Software, Inc. San Diego, USA). For statistical comparisons, one-way ANOVA multiple comparison (Kruskal-Wallis test, nonparametric test), unpaired t test (non-parametric Mann-Whitney test), or Fisher’s exact test was used. Statistical significance of survival was assessed with a Log-rank (Mantel-Cox) test. P values < 0.05 were considered significant. For the TMA-2 human adenocarcinoma cohort all statistics were performed in R. The cut-off for binarisation of SOX9 was obtained by a published method that has a greater power and lower bias than logrank optimisation in large data sets [31]. The proportional hazards assumption was tested by examination of log-log plots.

## Results

### The correlation between SOX9 expression and NSCLC staging is histopathology-selective

To evaluate SOX9 expression in NSCLC histotypes according to their stages, we performed IHC analyses of TMAs encompassing 28 human lung SCCs, 13 human lung ASCs and 25 human lung PACs. SOX9 expression was not seen in normal lung tissues but observed in the tumour regions in most of all samples (SCC: 96%, ASC: 100%, PAC: 88%). This significantly contrasts with the expression of SOX2, a well-established genetic driver of SCC, which was more commonly detected in SCCs and ASCs than PACs (SCC: 85%, ASC: 100%, PAC: 8%) (Figure 1A). The SOX9 protein was mainly located in the nucleoplasm, but cytoplasmic/membranous expression was also observed, particularly in PACs (Figure 1A-B).

**Figure 1.**
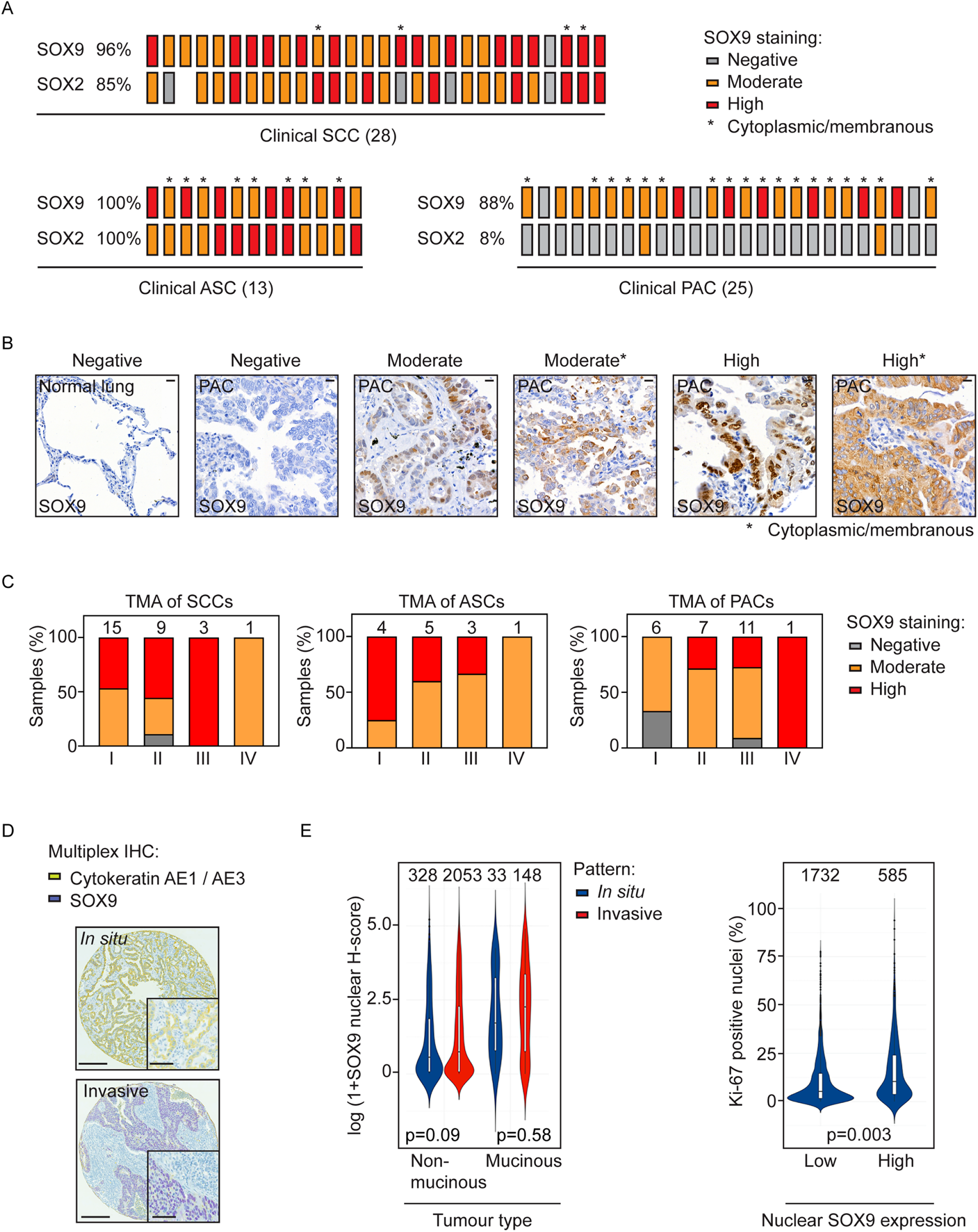
Histopathology-selective correlation between SOX9 expression and NSCLC progression. (A) Schematic summary of SOX9 and SOX2 expression, evaluated by visual scoring, in TMA panels representing clinical SCCs, ASCs and PACs. (B) Representative images illustrating SOX9 expression in normal and malignant epithelial lung tissues. Scale bar: 20 μm. (C) Contingency bar plots show percentages of SOX9-negative, -moderate and -high cases in each histotype, and for NSCLC staging categories determined following American Joint Committee on Cancer (AJCC) criteria. Sample sizes are indicated above each bar. (D) Representative images of duplex-stained *in situ* or invasive adenocarcinoma tumour tissues in TMA-2. Yellow = cytokeratin, purple = SOX9, blue = haematoxylin. Cores are 1 mm across. Scale bars: 200 μm for original or 500 μm for magnified images. (E) Violin plots showing the relationship between SOX9 and mucinous phenotype and local invasiveness in stained tissue cores (left), and between SOX9 status and tumour cell proliferation quantified by nuclear Ki-67 expression (right).

Based on visual scoring, we assigned each sample to a ‘SOX9-negative’, ‘SOX9-moderate’ or ‘SOX9-high’ category and grouped them by their clinical stages (Figure 1B-C). Among the three NSCLC histotypes, PACs contained the most SOX9-negative or-moderate cases, particularly in the lowest stage tumours (6/6 cases in stage I and 5/7 cases in stage II), whereas in SCCs and ASCs absence of SOX9 was only detected in one sample and at least half of all stage I-II cases were SOX9-high. Thus, a positive correlation between SOX9 expression and clinical stage was detected in PACs, but not in ASCs or SCCs.

To more deeply explore the relationship between SOX9 and the phenotype of human lung adenocarcinoma, we quantified the expression of SOX9 expression in epithelial tumour cell nuclei in a large cohort of 970 resected primary lung adenocarcinomas in tissue microarrays (Figure 1D). Interestingly, nuclear SOX9 levels were higher in mucinous than non-mucinous tumours (p<0.001), and cores with invasive growth patterns showed a non-significant trend towards higher levels of SOX9 expression selectively in non-mucinous samples (p=0.09 in non-mucinous; p=0.58 in mucinous) (Fig 1D-E). Finally, corroborating previous studies [19–20, 32], epithelial tumour cell proliferation assessed by the percentage of tumour nuclei expressing Ki-67 was significantly related to SOX9 expression (p=0.003) (Fig 1E), and in analysis of patient survival time after surgery higher median levels of SOX9 related to relatively poor outcome (Cox model HR=1.2, p=0.044) (supplementary material, Figure S1).

### SOX9 is commonly expressed in advanced murine *Kras^G12D^;Lkb1^-/-^* tumours

We subsequently applied animal models to study the role of SOX9 in NSCLC histotype formation. As a first step, we examined SOX9 expression in tumours from *Kras^G12D^;Lkb1^-/-^* mice (KL mice characterised in [25], in this study named KLS^wt^). Most KLS^wt^ histotypes, including ASC, MAC and PAC, displayed high SOX9 expression, but its expression was fully absent or low in IACs which are small and represent the least aggressive lesion type with low proliferation and oncogenic signalling activities (Figure 2; supplementary material, Figure S2A). Therefore, SOX9 expression levels in murine NSCLC histotypes were comparable to those in clinical NSCLC, showing absent or low expression in IAC or *in situ* non-mucinous adenocarcinomas and wide expression across all advanced NSCLC histotypes.

**Figure 2.**
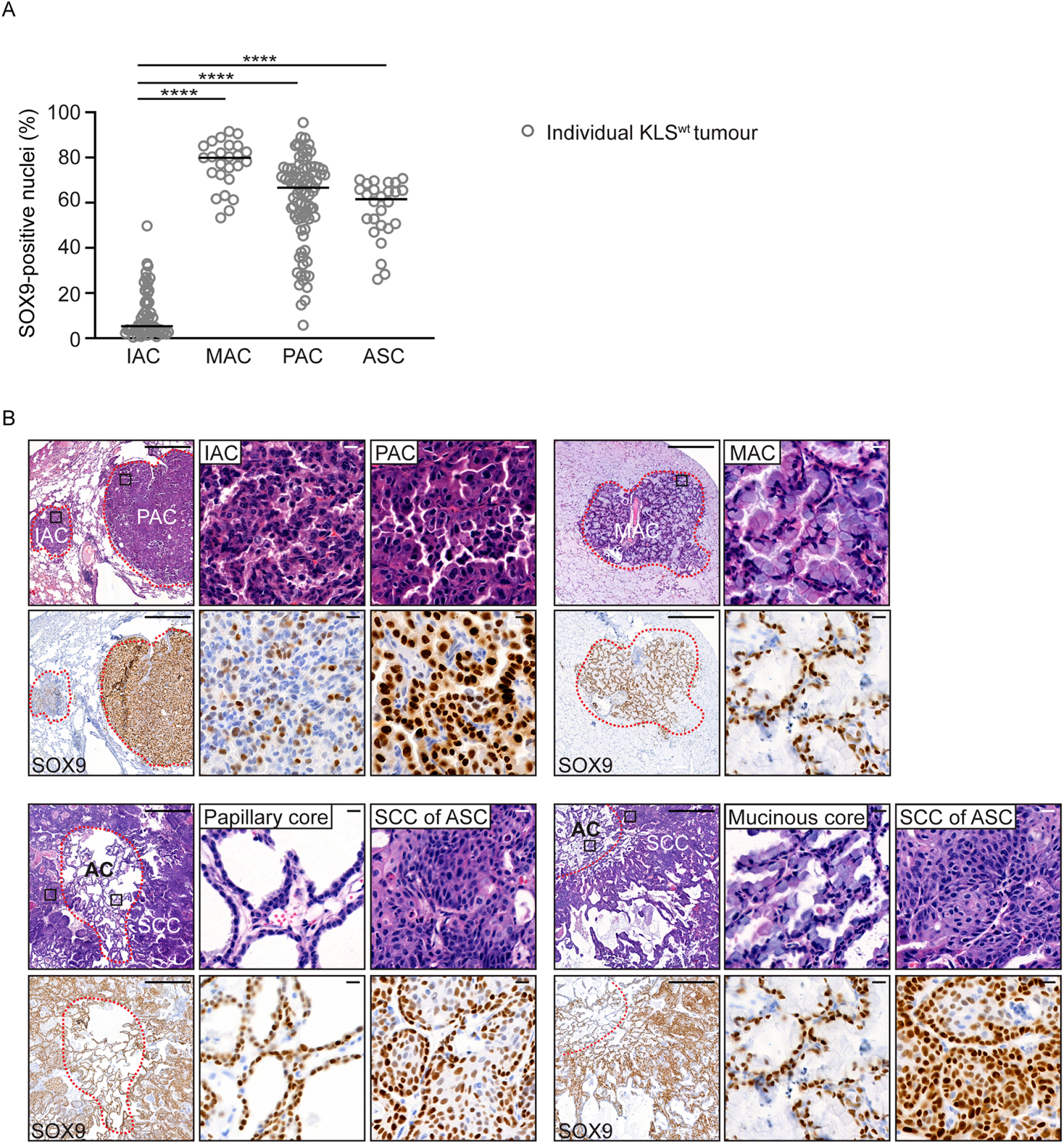
SOX9 expression is highest in advanced *Kras^G12D^;Lkb1^-/-^;SOX9^+/+^* murine NSCLC tumours. (A) Scatter dot plot showing SOX9 expression in individual tumours, grouped by histotype and calculated as percentages of SOX9-postive cells in total number of tumour cells, from six KLS^wt^-CC10 and eight KLS^wt^-SPC mice. Numbers above scatter plots indicate total tumour numbers. Statistical significance was assessed with One-way Anova (nonparametric Kruskal-Wallis test): ****p < 0.0001. (B) Representative H&E images and SOX9 stainings, illustrating SOX9 expression in each histotype. Areas depicted in higher magnification are outlined with black squares in the H&E images. Red dashed lines outline the whole tumour regions, or separate SCC and AC regions within an ASC. Scale bars: 500 μm for original or 10 μm for magnified images.

### SOX9 expression is essential for papillary structure formation in *Kras^G12D^;Lkb1^-/-^* murine tumours

To investigate the impact of *Sox9* loss on histotype formation, we established *Sox9*-deficient *Kras^G12D^;Lkb1^-/-^* cohorts, including KLS^het^ (losing one floxed *Sox9* allele) and KLS^null^ (losing two floxed alleles), following infection with either Ad5-CC10-Cre or Ad5-SPC-Cre viruses (Figure 3A).

**Figure 3.**
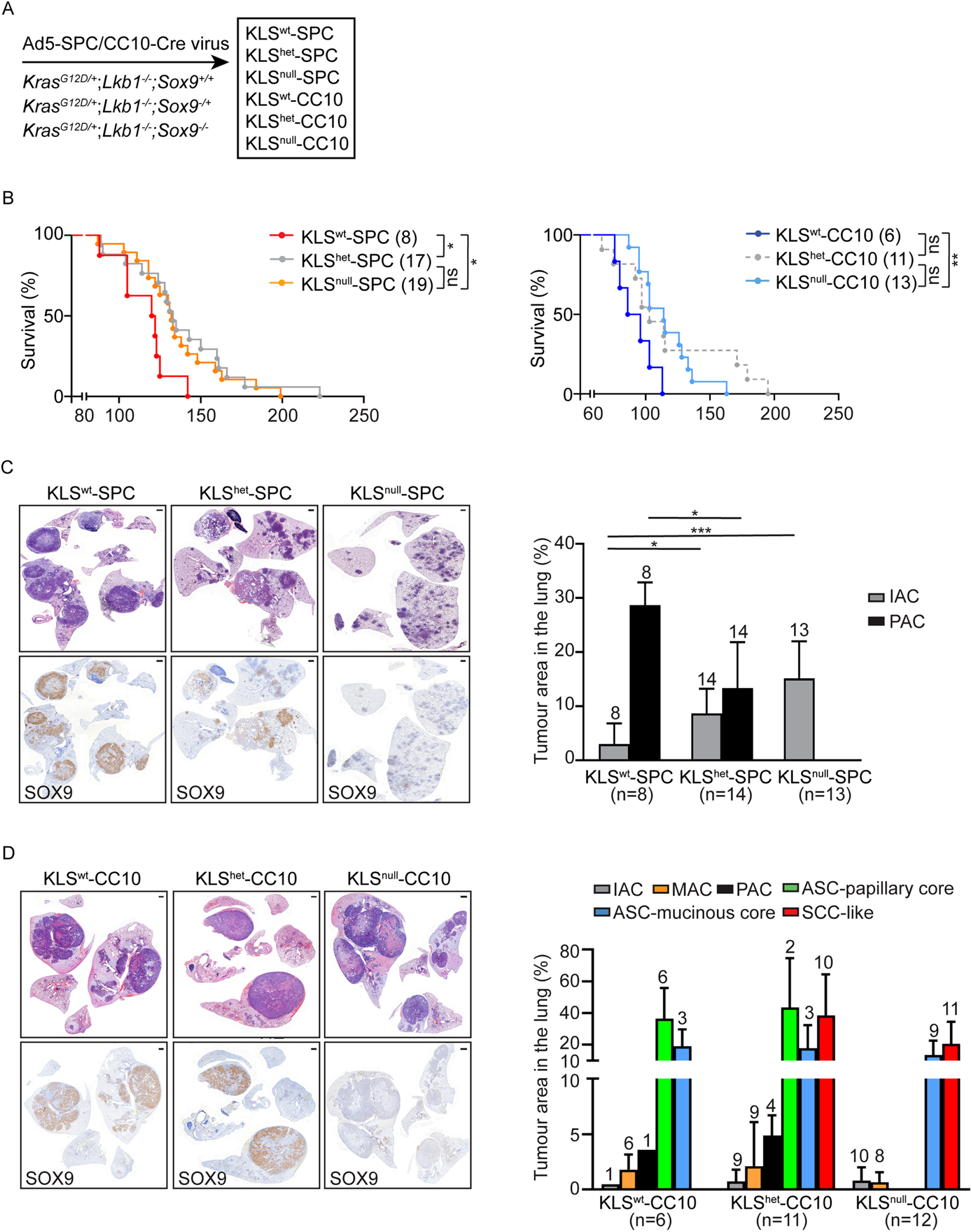
*Sox9* loss disrupts the formation of papillary histotype tissue in *Kras^G12D^;Lkb1^-/-^* NSCLC tumours. (A) Murine *Sox9* loss-of-function and control cohorts established for this study. (B) Kaplan-Meier survival curves for the indicated cohorts. Statistical significance was assessed with Log-rank (Mantel-Cox) test: *p < 0.05, **p < 0.01. (C) Left: representative images illustrating whole lung sections of individual mice from the indicated Ad5-SPC-Cre cohorts, depicting H&E staining and corresponding SOX9 expression. Scale bars: 1000 μm. Right: bar chart showing the histotype-selective tumour burden (% total histotype-specific tumour region per lung) in the indicated cohorts. Numbers of mice that carry the specified histotype are listed above each bar. In the KLS^null^-SPC cohort, only SOX9-negative tumours were plotted. Statistical significance was assessed with One-way Anova and non-parametric Kruskal-Wallis test: *p < 0.05, ***p < 0.001. (D) Left: representative images illustrating whole lung sections of individual mice from the indicated Ad5-CC10-Cre cohorts, depicting H&E staining and corresponding SOX9 expression. Scale bars: 1000 μm. Right: bar chart showing the histotype-selective tumour burden (% total histotype-specified tumour region per lung) in the indicated cohorts. Numbers of mice that carry the specified histotype are listed above each bar. In the KLS^null^-CC10 cohort, only SOX9-negative tumours were plotted.

Kaplan-Meier survival analysis showed significantly longer survival of KLS^het^-SPC and KLS^null^-SPC mice, as well as KLS^null^-CC10 mice, compared to the relative KLS^wt^ controls (Figure 3B). The increased life span of KLS^het^-SPC and KLS^null^-SPC cohorts corresponded to a reduced PAC burden (Figure 3C), and no papillary tissue structure was observed in SOX9-negative tumours in these mice (Figure 3C; supplementary material, Figure S2C). In Ad5-CC10-Cre cohorts, *Sox9* loss was associated with a shift in the major tumour burden from ASC to SCC tissue with large necrotic regions (Figure 3D). These SCC tumours were positive for the AC marker NKX2-1 and lacked SOX2 expression (supplementary material, Figure S2B), suggesting that these were derived from ASCs, and hence called SCC-like from here onwards. Interestingly, only the formation of ASC with papillary AC cores, but not those with mucinous AC cores, was disrupted (Figure 3D and supplementary material, Figure S2C). Additionally, PAC was the only AC subtype normally present in KLS^wt^-CC10 mice that was absent following *Sox9* loss (Figure 3D and supplementary material, Figure S2C).

Together, this data suggested that SOX9 expression was selectively required for the growth of papillary histotype tissue. In support of this conclusion, SOX9-positive tumours in both the *Sox9*^null^-SPC and -CC10 cohorts, explained by recombination escape known to arise in three-allelic compound mice [33], always showed a papillary tissue structure (supplementary material, Figure S3). As expected, based on the obligate role of LKB1 as a tumour suppressor in this model, we found no evidence for escape of *Lkb1* recombination (IHC staining; data not shown). Importantly, in both the CC10- and SPC-infected cohorts, KLS^het^ resembled KLS^null^ mice in terms of survival, histopathology spectra and tumour burden (Figure 3B-D; supplementary material, Figure S3C-D), together indicating that both heterozygous and homozygous *Sox9* loss selectively disrupted papillary NSCLC histotype formation.

### SOX9 expression is essential for the oncogenic transformation of cancer-initiating SPC+ cells

To understand why SOX9 selectively regulated the formation of papillary tissue structure, we studied the spatial expression of cell lineage markers and SOX9 in different tumour histotypes. In the Ad5-SPC-Cre cohorts, SPC, which marks AT2 progenitor cells and differentiated AT2 cells, was highly expressed in IACs, while SPC was absent or expressed at low levels in PACs (Figure 4A-B). Also in the Ad5-CC10-Cre cohorts, IACs were positive for SPC whereas PACs and papillary AC cores of ASCs showed negative to low SPC expression (Figure 4C-D; supplementary material, Figure S4A), indicating that also CC10-Cre induced the transformation of SPC+ cells. Other tumour histotypes found in Ad5-CC10-Cre cohorts that were not affected by *Sox9* loss (Figure 3D), particularly mucinous and squamous tissue, were always SPC-negative (Figure 4D and supplementary material, Figure S4A). Interestingly, we further found that SPC expression was mutually exclusive with SOX9 expression in PACs as well as papillary AC cores of larger ASC tumours, and marked tumour subregions that lacked prominent papillary structure (Figure 4B and D). The concomitant appearance of papillary structure together with SPC loss upon SOX9 expression, together with the absence of PAC and papillary AC cores following *Sox9* loss as shown in the section above, suggested that SOX9 was a driving force for papillary tumour progression from SPC+ progenitor cells (illustrated in Figure 4E).

**Figure 4.**
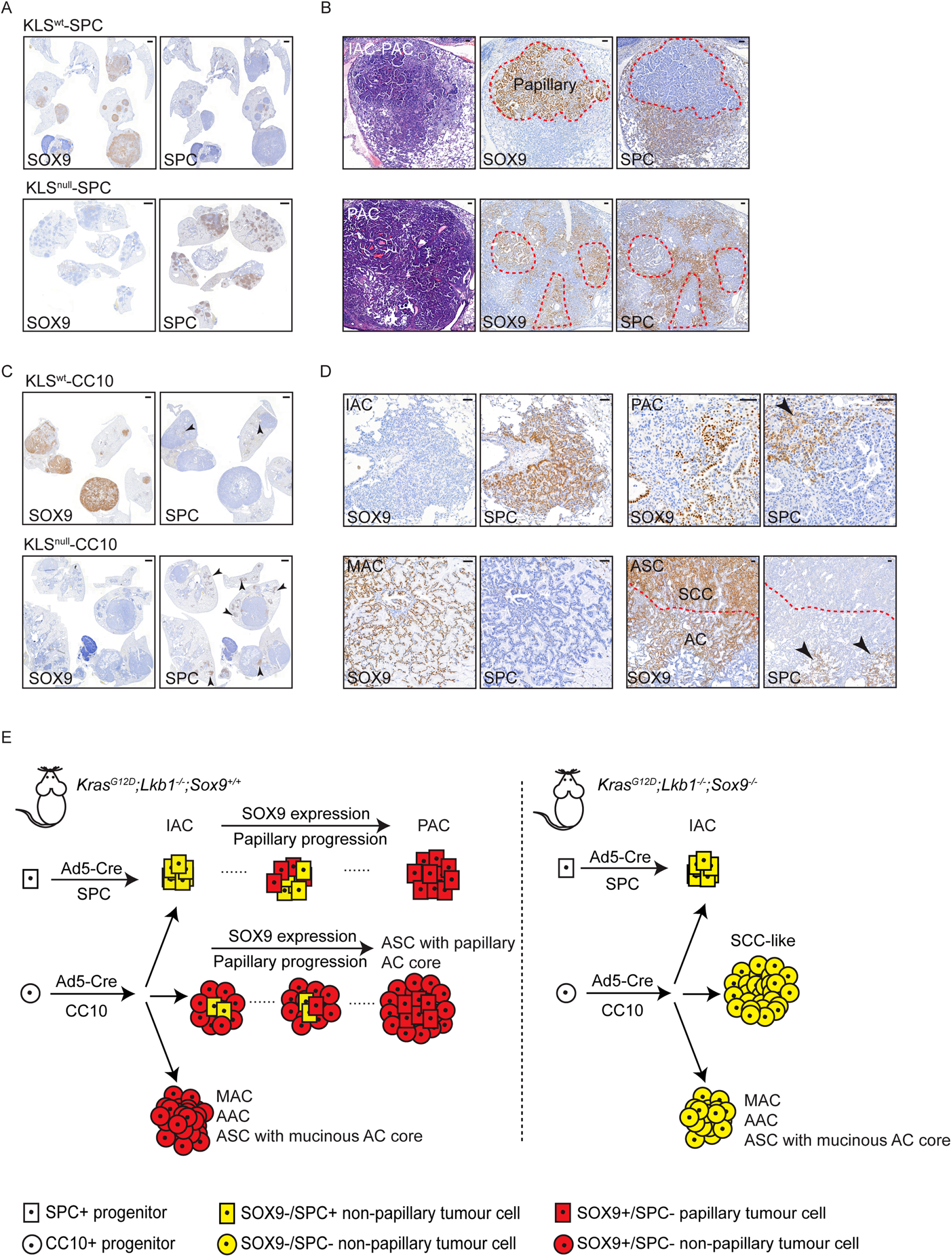
SOX9 expression drives the oncogenic transformation of SPC+ progenitors. (A) Representative images illustrating SOX9 and SPC stainings in the lung of a KLS^wt^-SPC and a KLS^null^-SPC mouse. More IACs were observed in the KLS^null^-SPC mouse, which all expressed SPC. Scale bars: 1000 μm. (B) Upper: representative images of a tumour from a KLS^null^-SPC mouse that had partially escaped *Sox9* recombination. The papillary area outlined by the red dashed line contained SOX9-positive regions that were SPC-negative; the remaining IAC-like region was SPC-positive. Lower: representative images of a PAC from a KLS^wt^ mouse, depicting mutual exclusive expression of SOX9 and SPC. Scale bars: 100 μm. (C) Representative images illustrating SOX9 and SPC stainings in the lung of a KLS^wt^-CC10 and a KLS^null^-CC10 mouse. Arrowheads point to SPC-positive tumour regions. Scale bars: 1000 μm. (D) Representative images showing spatial SOX9 and SPC expression in tumours of the indicated histotypes detected in KLS^wt^-CC10 mice. The red dashed line separates the SCC and AC regions within an ASC tumour. Arrowheads point to SOX9-negative, SPC-positive regions in a PAC and a papillary AC core region of an ASC. Scale bars: 100 μm. (E) Schematic explaining how SOX9 regulates murine NSCLC progression in a histotype-selective manner. Left: in KLS^wt^ mice, SPC+ AT2 cells, targeted by Ad5-SPC-Cre virus, give rise to SPC+ IACs. These IAC lesions gradually start to express SOX9 and this is essential for papillary tumour progression; the SPC+ identity is gradually lost when papillary tissue structure appears. Upon Ad5-CC10-cre infection, SPC+ IACs again develop, either from AT2 cells that are targeted by the CC10-cre virus or from trans-differentiated Clara or dual positive SPC+/CC10+ cells. SOX9 expression in papillary tissue corresponds with loss of SPC expression, giving rise to PACs or the papillary AC core of ASC. Lastly, transformation of CC10+ progenitor cells, which are mostly Clara cells, gives rise to MAC, AAC and ASC with mucinous AC core. Right: *Sox9* loss selectively disrupts papillary histotype tissue formation in which transformed cells exhibit progression from a SPC+ to a SPC-status.

### SOX9 expression differently regulates AC and SCC tissue metastases

Given the previous links between SOX9 expression and invasion, we next asked how *Sox9* loss affected the metastatic potential of NSCLC histotypes by analysing the mediastinal lymph nodes (supplementary material, Figure S4B-C). Metastases in KLS^wt^-SPC (observed in 3/8 KLS^wt^-SPC mice) were similar to those previously described [32] and represented AC histotype tissue with papillary structure (supplementary material, Figure S4C). Metastases that eventually developed in KLS^null^-SPC mice (4/19 mice, including 2 minimal metastases) were always SOX9-positive, suggesting that they had originated from PACs that had escaped *Sox9* recombination (Figure 5A and C). Also, the KLS^het^-SPC mice developed frequent metastasis (12/17 mice, including 6 minimal metastasis), which could however not be linked to SOX9 status due to the heterozygous SOX9 expression. Importantly, as previously reported [32], squamous metastases were never detected in KLS^wt^-CC10 mice, but observed in 2/11 KLS^het^-CC10 and 2/13 KLS^null^-CC10 mice (Figure 5B and D; supplementary material, Figure S4D). Three of the four mice carrying squamous metastases succumbed to their disease relatively early and within the survival range of control KLS^wt^ mice. This analysis therefore suggested an overall reduction in AC metastases following *Sox9* loss, linked to a block in PAC progression, mirrored by a notable increase in SCC-like metastases following *Sox9* loss in murine NSCLC.

**Figure 5.**
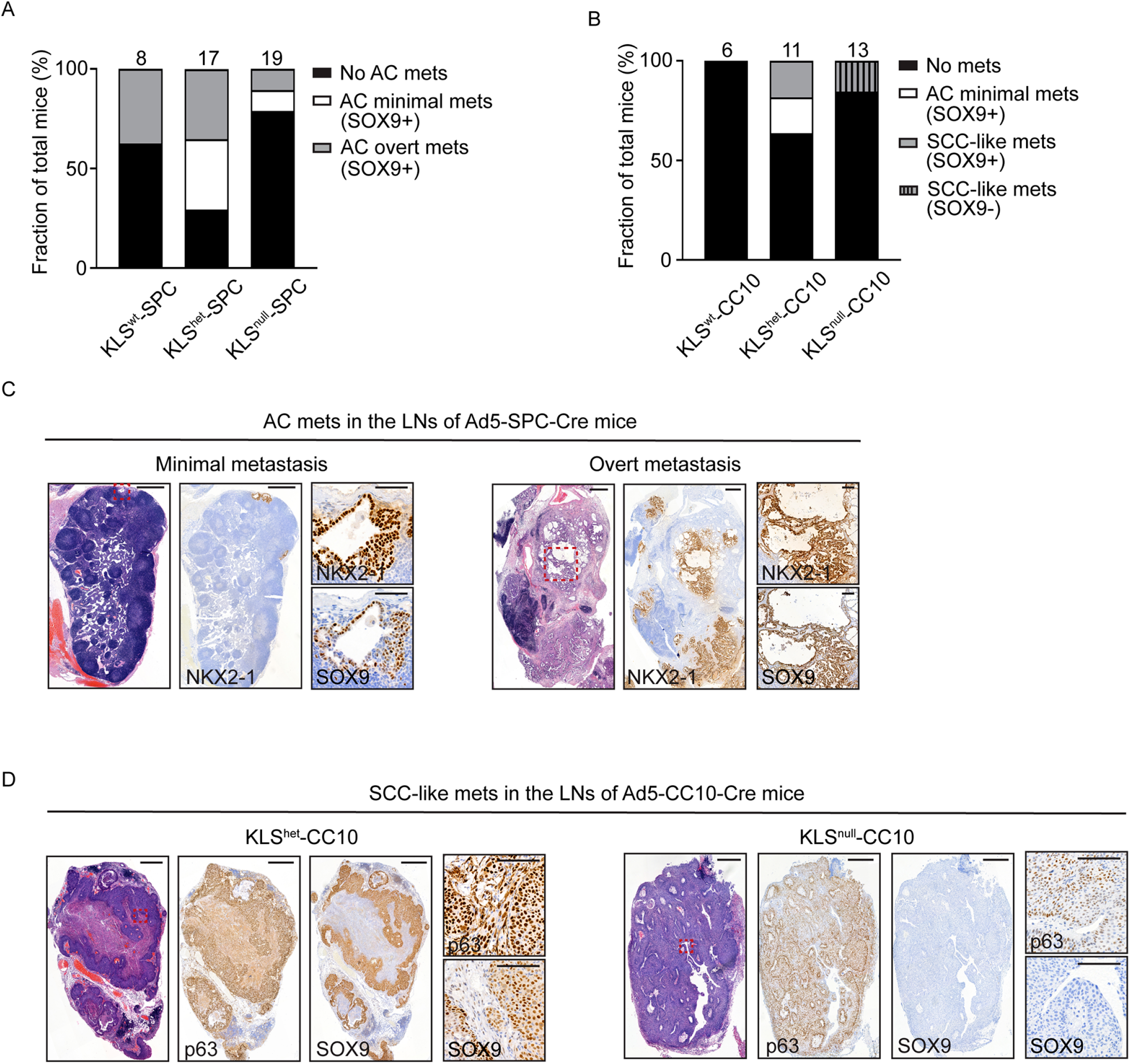
SOX9 expression differently regulates AC and SCC metastases. (A) Bar chart illustrating the proportion of total animals exhibiting no metastasis, minimal metastasis, or overt metastasis in the mediastinal lymph nodes (LNs) in the indicated Ad5-SPC-Cre cohorts. Total numbers of mice are listed above each bar. Metastases were all SOX9-positive, and all were adenocarcinoma. (B) Bar chart illustrating the proportion of total animals exhibiting no, minimal, or overt metastasis in the mediastinal LNs, subgrouped by SOX9 status, in the indicated Ad5-CC10-Cre cohorts. Total numbers of mice are listed above the bars. (C) Representative images illustrating minimal metastasis (only a small LN region infiltrated with tumour cells) and overt metastasis (LN structure severely disrupted) in Ad5-SPC-Cre cohorts; NKX2-1 indicates AC histotype tissue. Areas depicted in higher magnification are outlined with red dashed squares. Scale bars: 500 μm for original images or 100 μm for magnified images. (D) Representative images illustrating SCC-like metastases in Ad5-CC10-Cre cohorts; p63 indicates SCC histotype tissue. Areas depicted in higher magnification are outlined with red dashed squares. Scale bars: 500 μm for original images or 100 μm for magnified images.

As SOX9 is known to regulate extracellular matrix (ECM) deposition, we next examined ECM features following Sox9 loss by staining of collagen IV (COLIV), a central component of the tumour basement membrane. COLIV was expressed in both IAC and PAC, but only in the latter it formed a linear structure, which was also observed in papillary SOX9-expressing tissue at metastatic sites (Figure 6A-B). This suggested that a precise alignment of COLIV, regulated by SOX9 expression, might be required for the actual growth and later invasiveness of PACs. On the other hand, in *Sox9^wt^* ASCs, COLIV formed continuous boundaries around squamous tumour cell pockets (Figure 6C), while *Sox9^null^* SCC-like tumours and the SCC regions of ASCs with mucinous AC cores showed thinner parallel COLIV fibres as well as granular and focal depositions (Figure 6D). A similar alteration in COLIV rearrangement was also observed in the SCC-like LN metastases (Figure 6D). Taken together, in murine PACs, Sox9 loss reduced primary papillary tumour progression and consequently their invasion, possibly related to an alteration in ECM deposition, while in squamous histotype the altered COLIV deposition following *Sox9* loss in fact associated with an increase in tumour invasiveness and metastasis (illustrated in Figure 6E).

**Figure 6.**
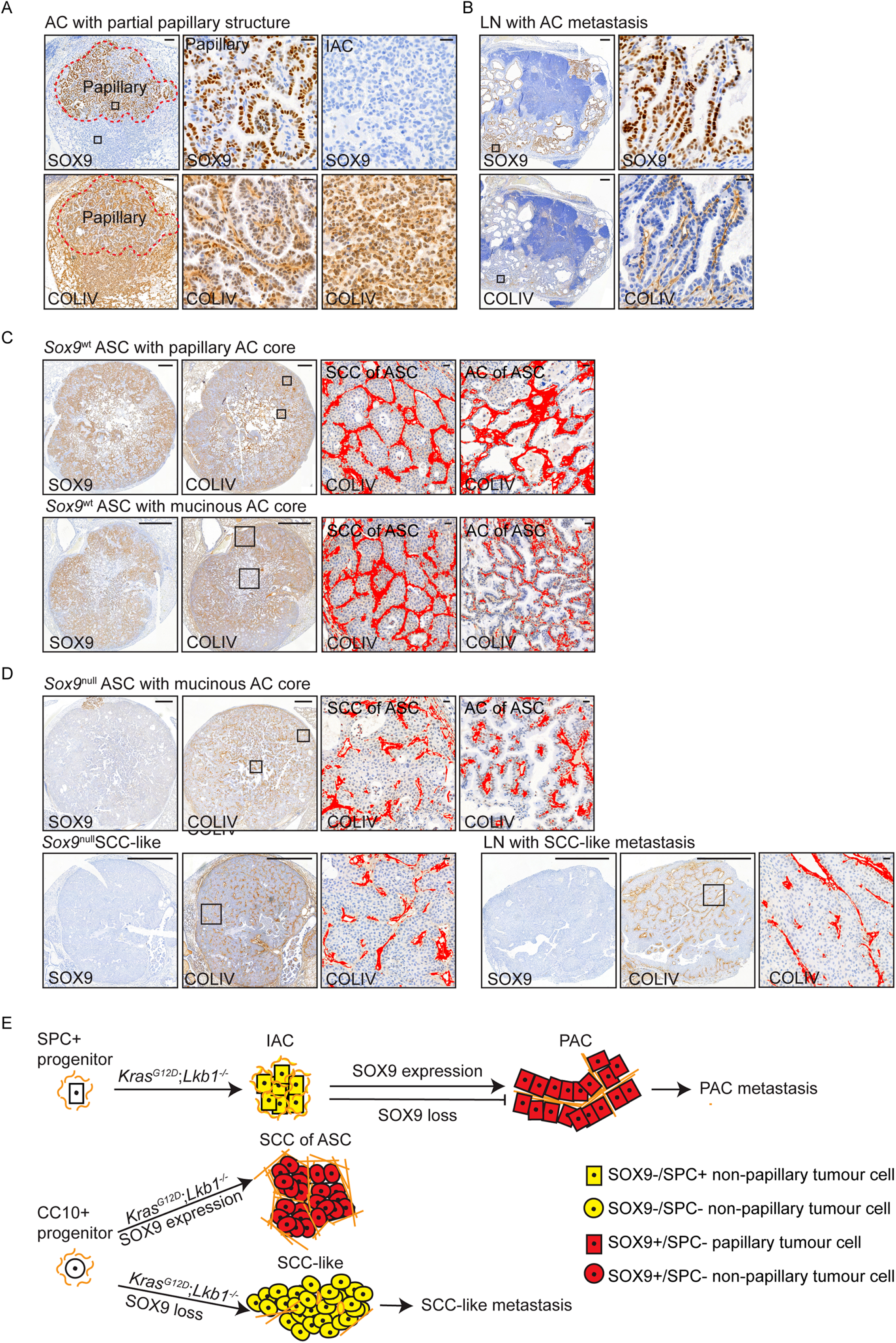
Loss of SOX9 affects collagen IV deposition in a histotype-selective manner. (A) Representative images showing SOX9 and collagen IV (COLIV) expression in a KLS^null^-SPC tumour with both PAC and IAC features. The red dashed line marks the papillary region. Squares mark the areas depicted in higher magnification. Scale bars: 100 μm for original images or 20 μm for magnified images. (B) Representative images showing SOX9 and COLIV expression in a LN metastasis with PAC features. Squares mark the areas depicted in higher magnification. Scale bars: 100 μm for original images or 20 μm for magnified images. (C) Representative images depicting the COLIV arrangement in a *Sox9^wt^*ASC with papillary AC core and a *Sox9^wt^ASC* with mucinous AC core. Squares mark the areas depicted in higher magnification. Scale bars: 1000 μm for original images or 20 μm for magnified images. (D) Representative images depicting the COLIV arrangement in a *Sox9^null^ASC* with mucinous AC core, a *Sox9^null^* SCC-like tumour, and a *Sox9^null^* SCC metastasis in the mediastinal LN (p63 staining of the same LN metastasis is shown in Fig. 5D). Altered COLIV deposition is detected selectively in SCC of ASC with mucinous AC core and SCC-like tissue. Squares mark the areas depicted in higher magnification. Scale bars: 1000 μm for original images or 20 μm for magnified images. (E) Schematic illustrating how SOX9 may regulate histotype-specific metastasis by influencing ECM (marked by COLIV) secretion/deposition. Linearised COLIV backbones form during IAC to PAC progression, which requires SOX9 expression. *Sox9* loss disrupts PAC formation, and hence their subsequent metastasis. To the contrary, while COLIV fibres form continuous boundaries around squamous tumour cell pockets in *Sox9*^wt^ SCC of ASC that may confine tumour invasiveness, they are re-oriented, or become granular/discontinuous following *Sox9* loss, in SCC-like tumours or SCCs of ASCs with mucinous AC cores. This associates with an increase in the incidence of SCC-like metastasis, and these metastases similarly show altered COLIV deposition.

## Discussion

Re-expression of the lung developmental transcription factor SOX9 specifically in malignant lung tissues, in conjunction with the reported positive correlation of SOX9 expression levels with clinical NSCLC staging, renders SOX9 a good candidate for prognostic and therapeutic interrogation. We here studied the previously unexplored expression and role of SOX9 in NSCLC histotypes. We identified a trend towards elevated SOX9 expression in non-mucinous clinical invasive AC compared to *in situ* samples, as well as in murine PAC compared to IAC, with IAC constituting an early form of PAC. SOX9 was widely expressed across multiple invasive stage murine and clinical NSCLC histotypes, yet murine *Sox9* loss only affected papillary structure formation, not the initiation or progression of other histotype lesions. The overarching conclusion of our work is therefore that the role of SOX9 in NSCLC is to be interpreted in a histotype-selective manner.

The expression of *Kras^G12D^* combined with *Lkb1* loss in alveolar AT2 or bronchial Clara lung progenitor cells gives rise to a range of NSCLC histotype tumours. While the alveolar differentiation marker SPC was uniformly expressed in SOX9-IAC, its downregulation coincided with the appearance of papillary tissue structure, and SPC was completely absent in mature SOX9-expressing papillary tumour regions. A transition from alveolar dedifferentiation to gradual SOX9 expression was similarly seen following oncogenic KRAS expression in organoids [34], and corroborates data showing that ectopic expression of SOX9 in embryonic lungs prevented AT2 cell differentiation [12,13], overall indicating that SOX9 expression intrinsically associates with AT2 lineage exit. Use of a conditional *Sox9* null model to address the role of SOX9 in NSCLC progression showed that *Sox9* loss disrupted the formation of PAC and papillary AC cores of ASC lesions, but not IAC or histotypes derived from bronchial progenitors. This therefore suggests that SOX9 expression is specifically required to drive the progression of lesions developed from a SPC+ cell of origin, leading to papillary NSCLC formation. Precisely how SOX9 drives AT2 lineage exit remains unclear, and an intriguing future direction would hence be to identify the lineage-specific SOX9 transcriptional network, and particularly its interplay with transcription factors known to regulate alveolar differentiation, such as FOXA2 [35–37]. Our study thus reveals a role of SOX9 in NSCLC progression that is selectively linked to the SPC+/AT2 lineage of cancer-initiating lung progenitor cells.

We showed that *Sox9* loss decreased AC metastasis, which is consistent with a reported prometastatic role of SOX9 in multiple cancer types [38–40]. But surprisingly, the incidence of SCC metastasis increased following *Sox9* loss. One mechanism in which SOX9 is involved in tumour metastasis is through regulating the expression and deposition of ECM components, including multiple collagens and laminins [14]. Our findings importantly suggest that SOX9-regulated ECM re-arrangements affect tumour invasiveness in different and microenvironmental context-dependent manners. We identified differential COLIV deposition, which marks tumour basement membranes, across NSCLC histotypes. Linear COLIV was observed in SOX9-expressing primary PACs and papillary metastatic LNs, but not in nonmetastatic SOX9-negative IACs, suggesting that it is part of a supportive structure for papillary growth and metastasis upon SOX9 expression. On the other hand, in the SOX9-expressing squamous regions COLIV formed pocket structures surrounding the tumour islets, and these became discontinuous and granular in SOX-negative squamous region. Similar COLIV discontinuity has been previously linked to the progression of oral SCC [41], suggesting that disrupted squamous pocket structure is commonly associated with increased squamous invasiveness. Our study therefore depicts functional heterogeneity of SOX9 in NSCLC metastasis, namely a requirement for SOX9-associated matrix deposition in papillary histotype metastasis, but an opposing metastasis-suppressing function in squamous histotype tissue.

Analysis of *Sox9* heterozygous null mice showed that loss of one *Sox9* allele already disrupted papillary tissue formation. While *Sox9* haploinsufficiency has not been previously described in cancer studies, loss of one *Sox9* allele has been associated with defects in beta-cell development during murine pancreas organogenesis [42], as well as with defects in murine chondrocyte differentiation [43]. In humans, *Sox9* haploinsufficiency caused by *de novo* heterozygous *Sox9* mutation leads to a rare but severe form of skeletal dysplasia called campomelic dysplasia (CD) [44], and CD mutations can act in a dominant negative manner by disrupting the capacity of SOX9 to dimerise [45]. While our study demonstrates a similar allelic dosage effect of *Sox9* in regulating NSCLC progression and metastasis, further studies are needed to understand whether this is associated with defective SOX9 dimerisation, and which are the target genes that essentially regulate histotype-selective effects during NSCLC progression.

Targeting transcription factors in cancer has been a long-standing challenge, yet the tumour cell-specific SOX9 expression renders it an attractive therapeutic target in NSCLC. Our results indicate that the inhibition of SOX9 could hamper the progression of early-stage ACs with SPC+/AT2 identity. Further study should address the effect of *Sox9* loss subsequent to PAC formation, for example using a flippase-FRT and Cre-loxP dual-recombinase system [46], to learn whether SOX9 is similarly required for the progression or metastasis of already established PACs. If so, then this would provide additional support favouring development of SOX9 inhibitors to treat PAC progression. Small molecules that disrupt the protein-interaction function of the SOX18 vascular development regulator have recently become available [47]. A similar strategy could possibly be applied to identify small molecules that disrupt SOX9 dimerisation or its binding to other lineage-specific transcription factors. However, while our study implies SOX9 as a potential anti-cancer target in papillary histotype NSCLC, *Sox9* loss also associated with enhanced squamous tissue metastasis, emphasising that therapeutic considerations ought to consider the tumour’s histotype context and cannot simply adopt SOX9 positivity as a biomarker for response.

In conclusion, we demonstrate opposing requirements for SOX9 in different NSCLC histotypes, and shed light on a pleiotropic SOX9 function that links to different tumour progenitor cells and extracellular microenvironmental features. This provides a strong motivation to evaluate the prognostic value and therapeutic application of SOX9 in the context of NSCLC histopathology types.

## Acknowledgements

We thank the Digital and Molecular Pathology Unit supported by the University of Helsinki and Biocenter Finland, and the Laboratory Animal Centre for husbandry support. We thank Zhangyi He for performing statistical analysis on the TMA-2 dataset. Katja Välimäki is thanked for material support, Ashwini Nagaraj for advice, and Iris Lähdeniemi for reviewing the manuscript. Samples/data used for the research were obtained from the Helsinki Biobank, and we warmly thank all patients for consenting to archive their samples in the Biobank. Research was supported by the University of Helsinki Doctoral Programme in Biomedicine (JB); The Finnish Medical Foundation (MIM); MRC Toxicology Unit core funding (JLQ); Innovative Medicines Initiative Joint Undertaking grant agreement no. 115188, the resources of which are composed of a financial contribution from the European Union’s Seventh Framework Programme (FP7/2007-2013) and EFPIA companies’ in-kind contribution (EWV); and the Academy of Finland grant 307111 (EWV).

## Author contributions

JB, KN and EWV designed the study and planned the experiments. KN established the animal cohorts and started the experiments. JB conducted further experiments and performed histopathology-selective analyses. AT performed immunohistochemistry and quantification on the large clinical adenocarcinoma cohort. AH and NML assisted with animal health checks, genotyping, and tissue processing. AH performed image scanning. KS provided histopathological review of the murine tumours. MIM provided the grade information of clinical NSCLC samples. JLQ supervised the large clinical adenocarcinoma cohort analysis. JB and EWV wrote the manuscript, and EWV provided supervision.

## Data availability statement

The data that support the findings of this study, including murine tumour and TMA-1 analyses, are available as supplementary materials, Table S2–3. Data generated with TMA-2 is accessible by contacting the authors.

---

**Figure S1.**
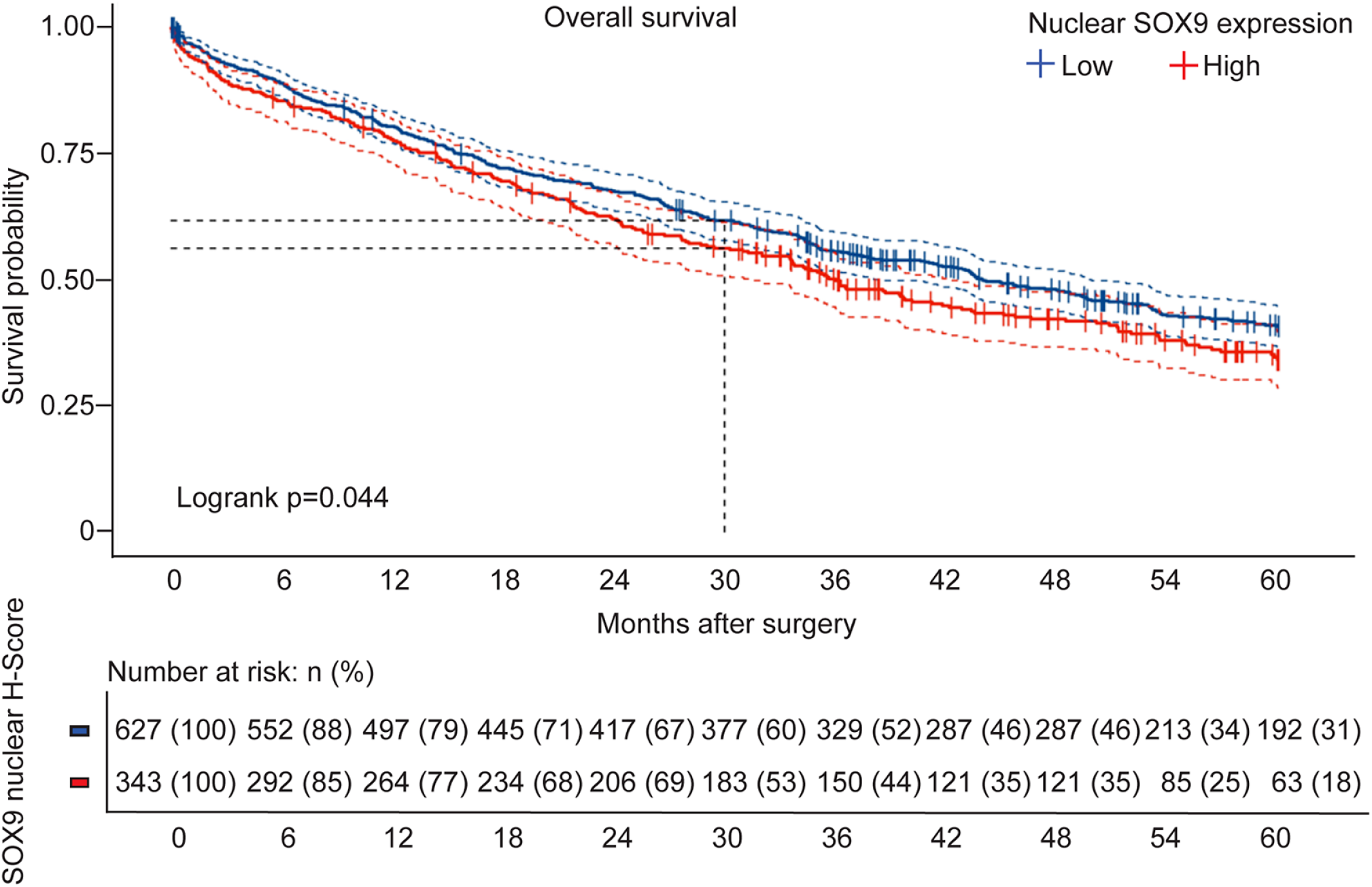
Cox proportional hazards model of patient survival after surgery related to SOX9 expression. Higher median nuclear SOX9 expression is significantly associated with relatively poor overall survival in patients with lung AC after surgery.

**Figure S2.**
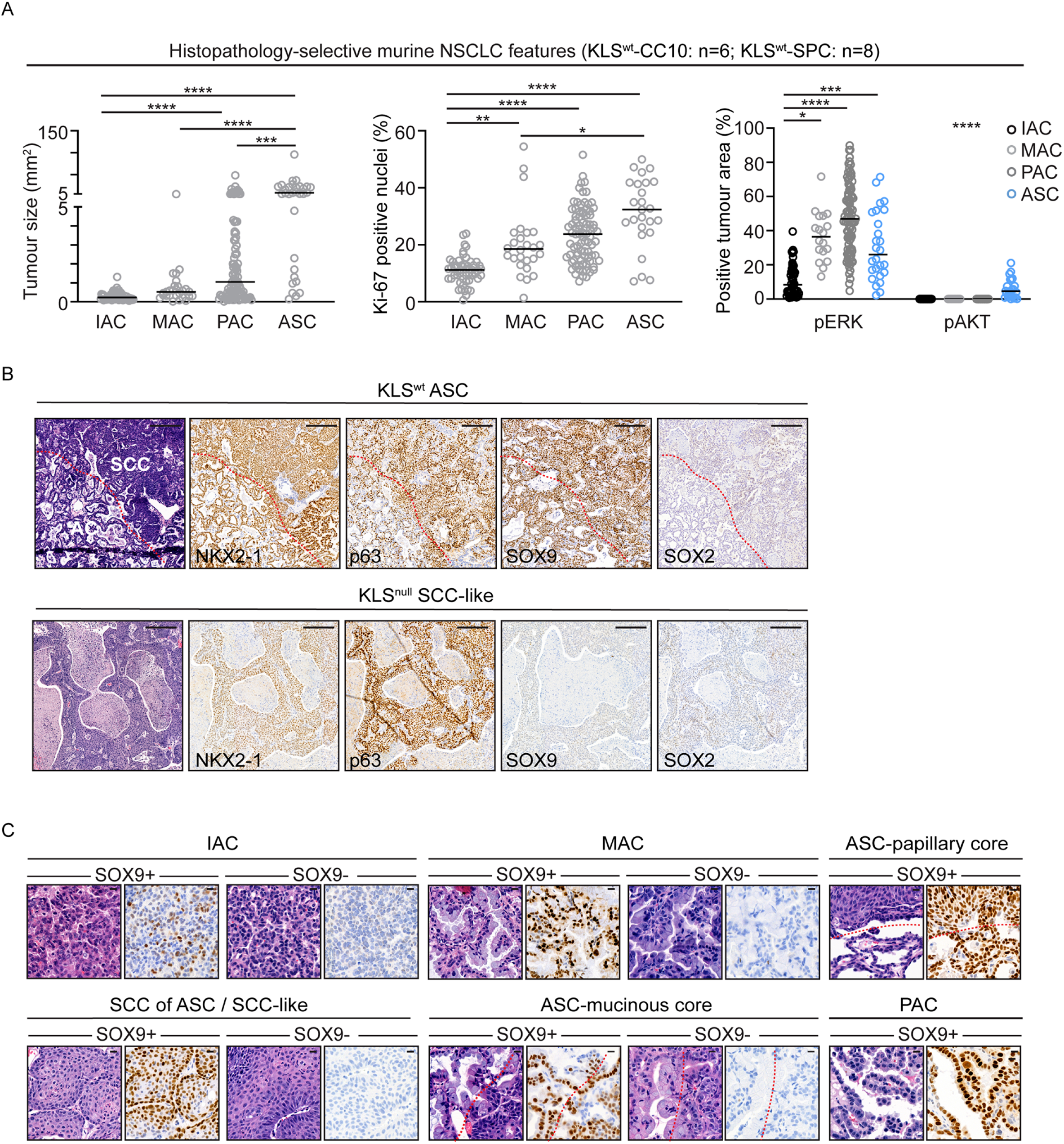
Altered histopathology and tumour burden following heterozygous or homozygous *Sox9* loss. (A) Three scatter dot plots summarising the size (left), proliferation rate (centre) and MAPK/AKT oncogenic signalling activities (right) in murine NSCLC histotypes. Stainings were performed on serial tissue sections and analysed by histopathology-guided analyses of individual tumours. Statistical significance was assessed with One-way Anova (non-parametric Kruskal-Wallis test): *p < 0.05, **p < 0.01, ***p < 0.001, ****p < 0.0001. (B) Representative images depicting marker stainings for murine ASC and SCC-like histotypes. NKX2-1: AC marker; p63: SCC marker. Scale bars: 100 μm. (C) Representative images depicting murine NSCLC histotype tissues from KLS^wt^ and KLS^null^ cohorts that express or lack detectable SOX9. Scale bars: 10 μm. The red dashed line separates the AC and SCC regions within an ASC. PACs or ASCs with papillary AC cores were not detected in the absence of SOX9 expression.

**Figure S3.**
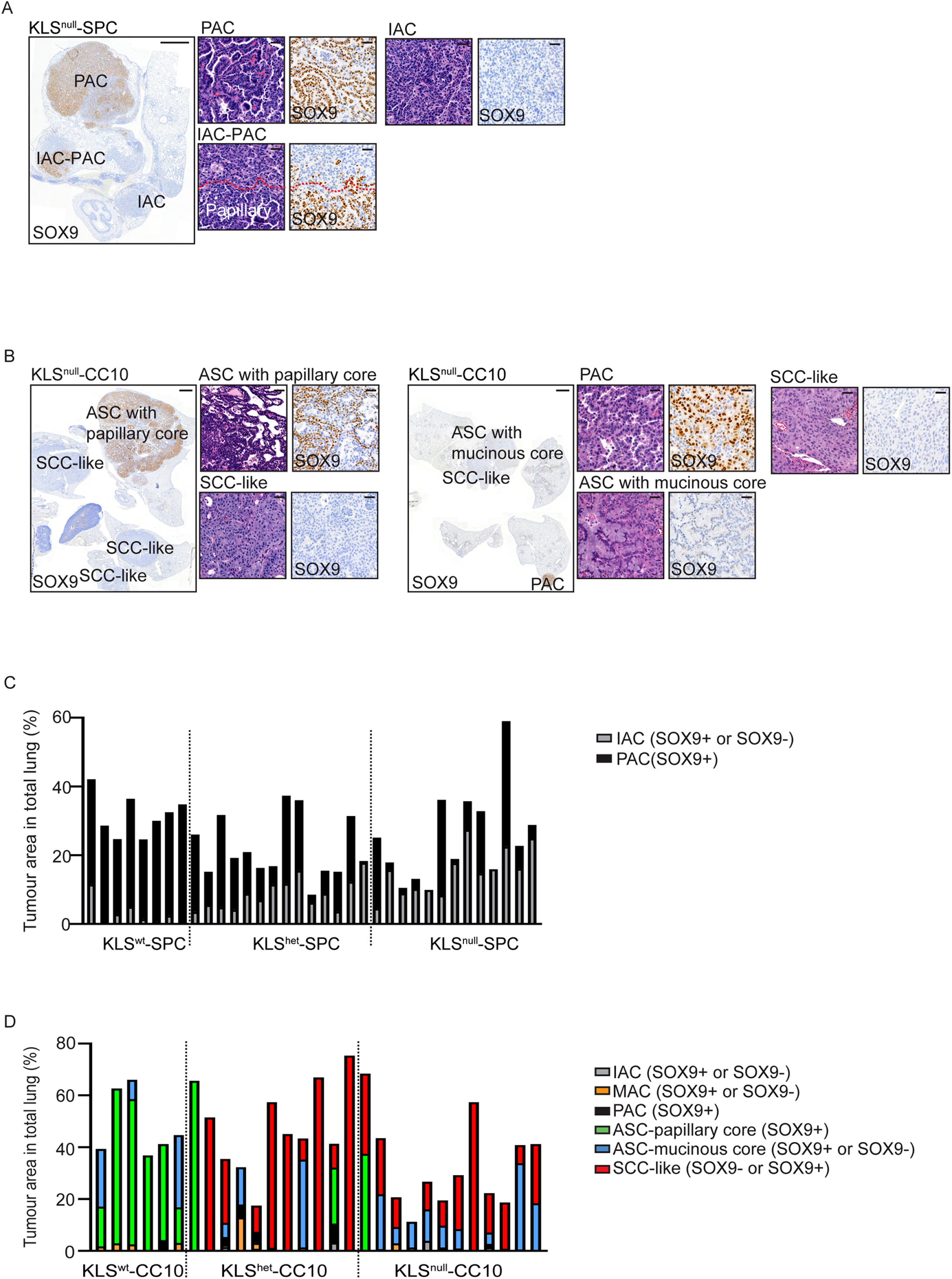
Papillary histotype formation as a result of escaping Sox9 recombination in KLS^null^ cohorts. (A) Representative images of PAC tumours that developed following escape from *Sox9* recombination in a KLS^null^-SPC mouse. The red dashed line separates the papillary histotype region from its precursor IAC lesion. Scale bars: 1000 μm for full images and 20 μm for magnified images. (B) Representative images showing the appearance of an ASC with papillary AC core (left) and a PAC (right) that developed following escape from *Sox9* recombination in two KLS^null^-CC10 mice. Scale bars: 1000 μm for full images and 20 μm for magnified images. (C) Contingency bar plot showing histotype-specific tumour burdens (% area of each histotype perlung) in eight KLS^wt^-SPC, 14 KLS^het^-SPC, and 13 KLS^null^-SPC mice. All tumours, including those that escaped Sox9 recombination, are plotted. PACs found in KLS^null^-SPC mice were always SOX9-positive. (D) Contingency bar plot showing histotype-specific tumour burdens (% area of each histotype per lung) in six KLS^wt^-CC10, 11 KLS^het^-CC10, and 12 KLS^null^-CC10 mice. All tumours, including those that escaped Sox9 recombination, were plotted. PACs and ASCs with papillary AC cores detected in KLS^null^-CC10 mice were always SOX9-positive.

**Figure S4.**
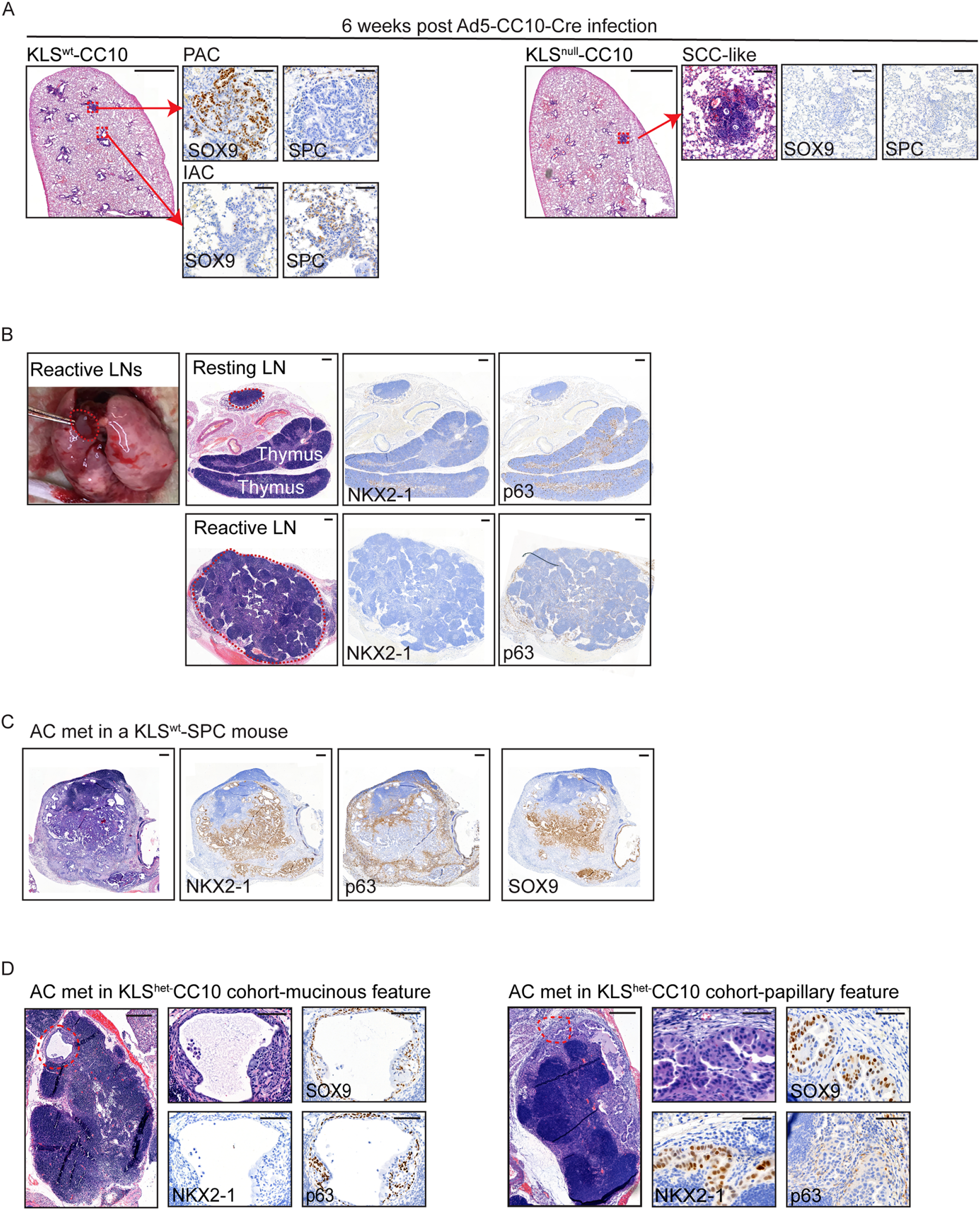
The histopathology-selective effect of SOX9 on the progression and metastasis of murine NSCLC tumours. (A) Left: representative images of a KLS^wt^ lung lobe collected six weeks post Ad5-CC10-Cre infection. One small lesion was SOX9+/SPC- and had papillary structure, while another was SOX9-/SPC+ with IAC histotype features. Right: representative images of a KLS^null^ lung lobe collected six weeks post Ad5-CC10-Cre infection, where the highlighted small lesion was SOX9-/SPC- and SCC-like. Scale bars: 1000 μm for the lung overview image and 100 μm for the magnified images. (B) Representative H&E images showing resting and reactive mediastinal LNs (outlined with a red dashed line) and the macroscopic location of LNs collected in this study. The tissue under the resting LN represents the thymus. No metastasis was found in resting and reactive LNs, confirmed by staining with the AC marker NKX2-1 and SCC marker p63. Scale bars: 200 μm. (C) Representative images illustrating AC metastasis in a KLS^wt^-SPC mouse. The metastatic tissue is NKX2-1+, SOX9+ and p63-in epithelial regions. Scale bars: 200 μm. (D) Representative images showing a mucinous minimal metastasis positive for the SCC marker p63, and an AC minimal metastasis positive for the AC marker NKX2-1 observed in KLS^het^-CC10 mice. SOX9 staining was positive in both cases. Outlined regions are represented in the magnifications. Scale bars: 500 μm for images showing entire LNs and 100 μm for magnifications.

**Table S1.** Primary antibodies used in the study (for TMA-1 and murine samples)

| **Antibody** | **Reference** | **Dilution** | **Validation method** |
| --- | --- | --- | --- |
| Rabbit anti-SOX9 | Millipore 5535 | 1:10000 | positive and negative reference tissues (*Kras*<sup>G12D</sup>;*Lkb1*<sup>-/-</sup>, *Kras*<sup>G12D</sup>;*Lkb1*<sup>-/-</sup>;*Sox9*<sup>-/-</sup> driven mouse lung tumours and normal murine epithelium) |
| Rabbit anti-Prosurfactant Protein C | Millipore 3786 | 1:2000 | positive and negative reference tissues (normal murine epithelium and *Kras*<sup>G12D</sup>;*Lkb1*<sup>-/-</sup> driven IAC tumours, *Kras*<sup>G12D</sup>;*Lkb1*<sup>-/-</sup>;*Sox9*<sup>-/-</sup> driven SCC-like tumours) |
| Rabbit anti-Collagen IV | Ab 6586 | 1:1500 | positive reference tissue (*Kras*<sup>G12D</sup>;*Lkb1*<sup>-/-</sup> ASC tumor ) |
| Goat anti-SOX2 | Santa Cruz Biotechnology sc 17320 | 1:200 | positive reference tissue (TMA of clinical NSCLC SCC sample) |
| Rabbit anti-NKX2-1 (TTF-1) | Ab 133638 | 1:2000 | positive reference tissue (*Kras*<sup>G12D</sup>;*Lkb1*<sup>-/-</sup> AC tumour) |
| Rabbit anti-p63 | Ab124762 | 1:10000 | positive reference tissue (*Kras*<sup>G12D</sup>;*Lkb1*<sup>-/-</sup>, ASC tumor) |
| Rabbit anti-pAKT (S473) | CST 4058 | 1:400 | PI3K/AKT inhibition (BEZ235) in murine NSCLC slice |
| Rabbit anti-p44/22 (Erk1/2) | CST 4370 | 1:1000 | MEK inhibition in murine NSCLC slice; blocking peptide (from CST); positive reference tissue (embryonic lung day 14) |
| Rabbit anti-Ki67 | Thermo Scientific RM-9106-S0 | 1:500 | Positive reference tissue (murine embryonic tissue, *Kras* lung tumors) |

For all the antibodies except anti-SPC (which did not require antigen retrieval), antigen retrieval was performed using 10mM sodium citrate pH 6.0. All antibodies except anti-SOX2 were incubated 1.5 h at room temperature while anti-SOX2 antibody was incubated overnight at 4°C.
CST = Cell Signaling Technology, Danvers, MA, USA
Ab = Abcam, Cambridge, UK

**Supplementary material, TableS2.**
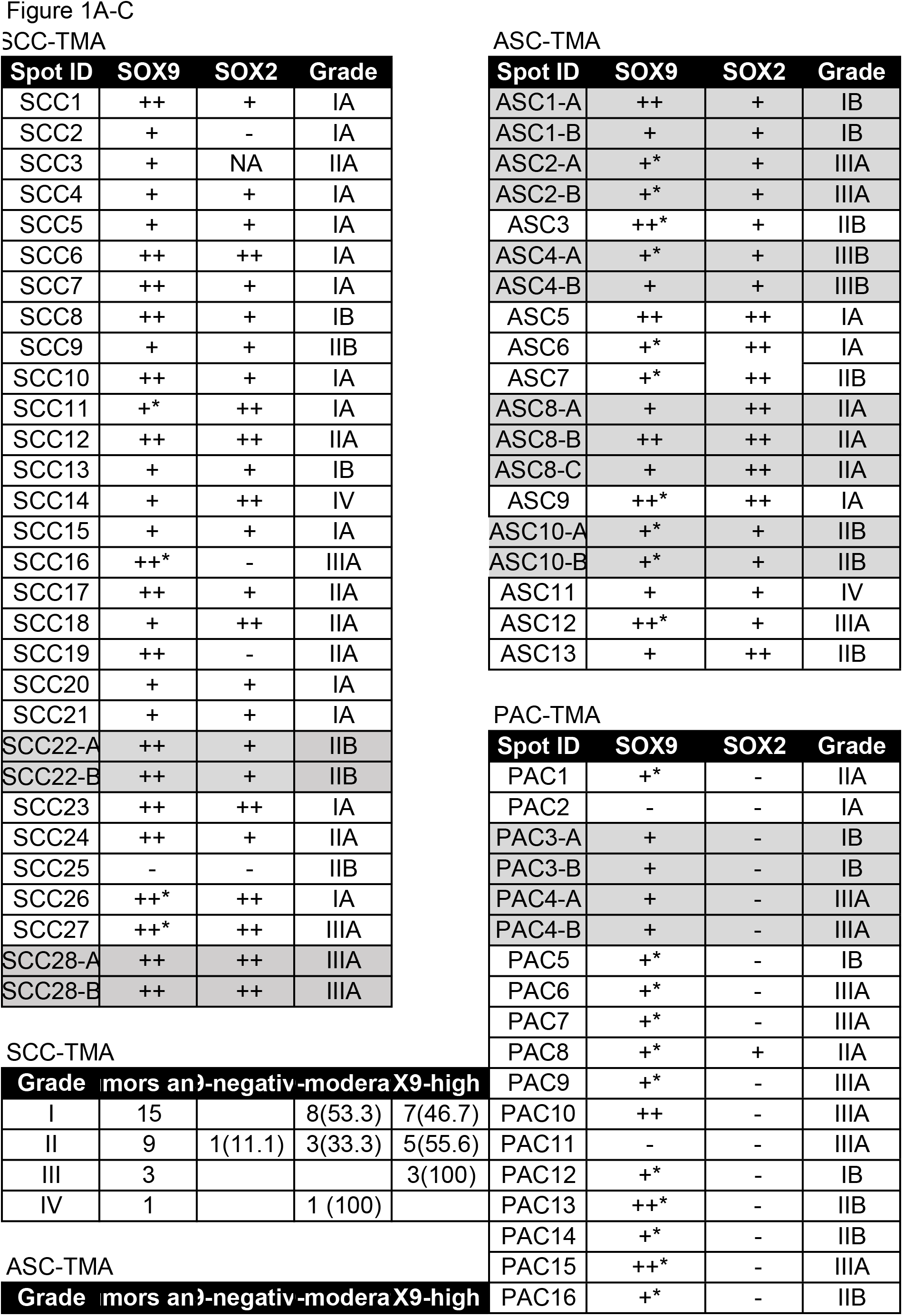
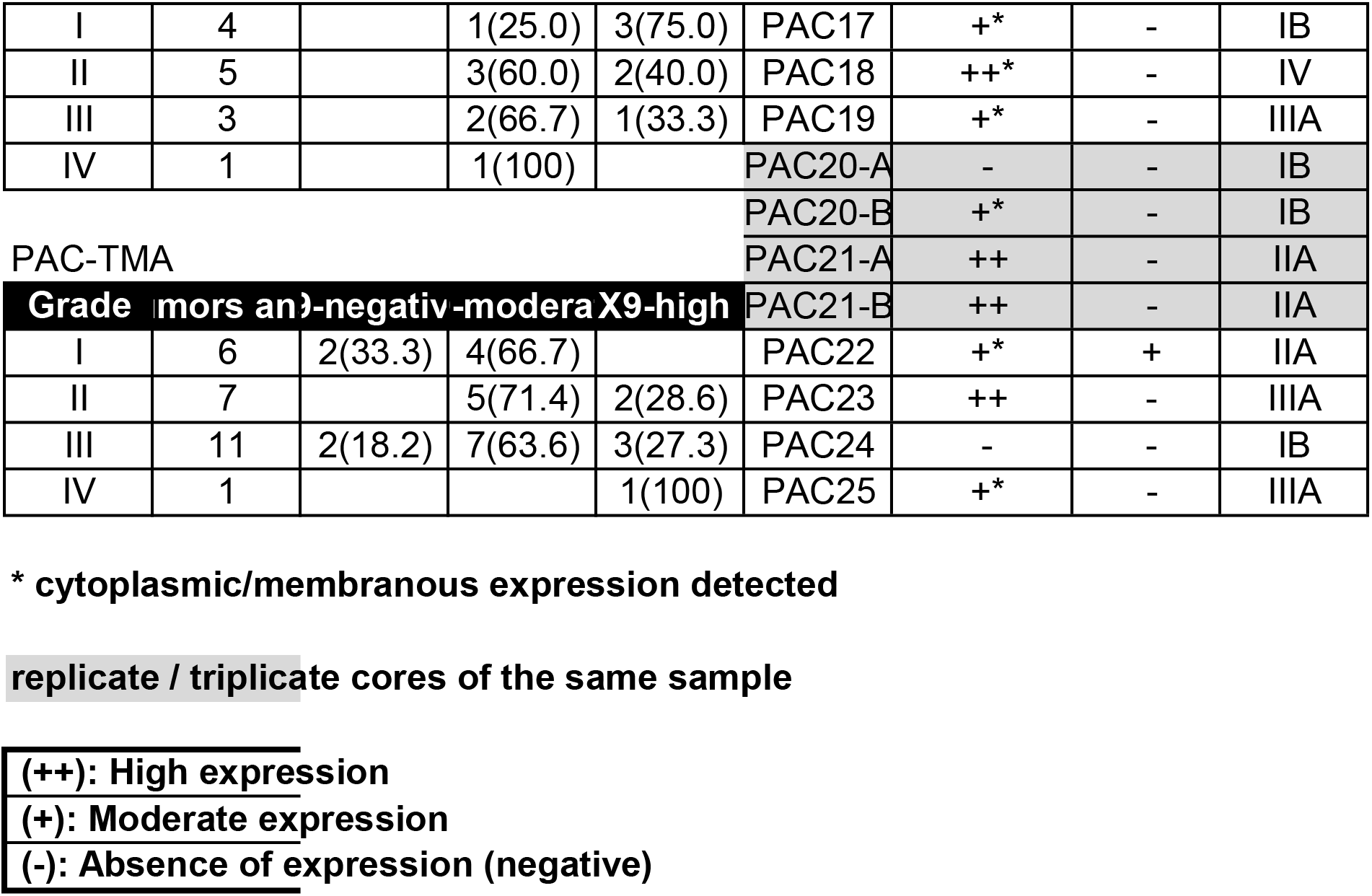
Human_NSCLC_TMA_analyses.

**Supplementary material, TableS3.** Murine_tumour_analyses.

| Tumour | Size(mm2) | %SOX9 | %Ki67 | %pERK | %pAKT |
| --- | --- | --- | --- | --- | --- |
| IAC-1 | 0.603 | 0.86 | 12.27 | 4.73 | 0 |
| IAC-2 | 0.475 | 24.84 | 14.41 | 1.58 | 0 |
| IAC-3 | 0.278 | 32.07 | 9.76 | 8.9 | 0 |
| IAC-4 | 0.364 | 15.49 | NA | 16.26 | 0 |
| IAC-5 | 0.167 | 24.84 | 10.09 | 2.16 | 0 |
| IAC-6 | 0.493 | 26.71 | 7.5 | 0.78 | 0 |
| IAC-7 | 0.273 | 9.15 | 6.7 | 1.34 | 0 |
| IAC-8 | 0.535 | 20.12 | NA | 4.34 | 0 |
| IAC-9 | 0.213 | 8.91 | 20.09 | 18.61 | 0 |
| IAC-10 | 0.404 | 5.08 | 8.11 | 2.68 | 0 |
| IAC-11 | 0.167 | 1.36 | 9.26 | 1.21 | 0 |
| IAC-12 | 0.168 | 5.26 | 4.33 | 24.58 | 0 |
| IAC-13 | 0.443 | 3.77 | NA | NA | 0 |
| IAC-14 | 0.404 | 1.6 | 15.51 | 14.89 | 0 |
| IAC-15 | 0.412 | 11.5 | 12.1 | 17.82 | 0 |
| IAC-16 | 0.267 | 5.1 | 14.06 | 13.92 | 0 |
| IAC-17 | 0.126 | 3.54 | 9.28 | 16.69 | 0 |
| IAC-18 | 0.239 | 2.91 | 9.36 | 2.98 | 0 |
| IAC-19 | 1.308 | 5.46 | 11.24 | 8.68 | 0 |
| IAC-20 | 0.134 | 2.77 | 13.04 | 14.31 | 0 |
| IAC-21 | 0.62 | 2.73 | 11.55 | 2.52 | 0 |
| IAC-22 | 0.188 | NA | NA | 8.07 | 0 |
| IAC-23 | 0.74 | 5.24 | 11.43 | 11.28 | 0 |
| IAC-24 | 0.256 | NA | NA | NA | NA |
| IAC-25 | 0.28 | NA | NA | NA | NA |
| IAC-26 | 0.458 | 15.87 | 5.79 | 15.87 | 0 |
| IAC-27 | 0.592 | 21.09 | 23.47 | 21.09 | 0 |
| IAC-28 | 0.468 | 26.93 | 23.94 | 26.93 | 0 |
| IAC-29 | 0.586 | 16.55 | 19.41 | 16.55 | 0 |
| IAC-30 | 0.168 | 4.54 | 9.26 | 4.54 | 0 |
| IAC-31 | 0.183 | 11.24 | 4.21 | 11.24 | 0 |
| IAC-32 | 0.478 | 15.64 | 8.11 | 15.64 | 0 |
| IAC-33 | 0.32 | 9.25 | 2.23 | 9.25 | 0 |
| IAC-34 | 0.381 | 6.79 | 0.59 | 6.79 | 0 |
| IAC-35 | 0.305 | 3.04 | 10.5 | 0.96 | 0 |
| IAC-36 | 0.186 | 2.59 | 9.3 | 3.916 | 0 |
| IAC-37 | 0.116 | 2.29 | 14.81 | 1.34 | 0 |
| IAC-38 | 0.105 | 6.16 | 12 | 4.9 | 0 |
| IAC-39 | 0.145 | 22.34 | 11.19 | 39.5 | 0 |
| IAC-40 | 0.134 | 3.81 | 12.6 | 28.68 | 0 |
| IAC-41 | 0.123 | 3.2 | 17.69 | 3.47 | 0 |
| IAC-42 | 0.09 | 10.26 | 14.07 | 6.89 | 0 |
| IAC-43 | 0.128 | 3.67 | 10.17 | 0.97 | 0 |
| IAC-44 | 0.089 | 10.43 | 14.02 | 2.03 | 0 |
| IAC-45 | 0.104 | 3.69 | 12.66 | 4.8 | 0 |
| IAC-46 | 0.42 | 2.4 | 12.77 | 1.8 | 0 |
| IAC-47 | 0.06 | 2.9 | 12.78 | 3.28 | 0 |
| IAC-48 | 0.149 | 33.03 | 11.35 | 2.22 | 0 |
| IAC-49 | 0.135 | 29.28 | 10.04 | 5.45 | 0 |
| IAC-50 | 0.111 | 49.81 | 8.06 | 15.58 | 0 |
| IAC-51 | 0.232 | NA | 12.76 | 0.01 | 0 |
| IAC-52 | 0.114 | 21.15 | 16.77 | 17.76 | 0 |
| IAC-53 | 0.269 | NA | NA | 2.93 | 0 |
| IAC-54 | 0.193 | 0.78 | NA | 8.75 | 0 |
| IAC-55 | 0.241 | 3.46 | NA | 3.51 | 0 |
| IAC-56 | 0.113 | NA | 11.51 | 19.57 | 0 |
| IAC-57 | 0.508 | 0.52 | 4.11 | 27.72 | 0 |
| IAC-58 | 0.085 | 5.83 | 7.17 | 27.69 | 0 |
| IAC-59 | 0.231 | 2.28 | 8.14 | 11.49 | 0 |
| IAC-60 | 0.092 | 1.12 | 3.71 | 20.62 | 0 |
| IAC-61 | 0.146 | 1.9 | 3.89 | 17.71 | 0 |
| IAC-62 | 0.101 | NA | NA | NA | NA |
| IAC-63 | 0.152 | 7.95 | 10.91 | 11.96 | 0 |
| MAC-1 | 0.017 | 61.35 | 25.64 | 49.96 | 0 |
| MAC-2 | 1.1 | 56.5 | 8.69 | 34.1 | 0 |
| MAC-3 | 1.7 | 63.2 | 15.28 | 40.6 | 0 |
| MAC-4 | 0.651 | 73.36 | 14.47 | 32.9 | 0 |
| MAC-5 | 0.451 | 82.55 | 22.28 | 51.37 | 0 |
| MAC-6 | 0.06 | 78.28 | 46.78 | NA | NA |
| MAC-7 | 0.206 | 81.07 | 9.04 | 48.8 | 0 |
| MAC-8 | 0.087 | 75.55 | 54.55 | 21.23 | 0 |
| MAC-9 | 1.5 | 90.59 | NA | NA | NA |
| MAC-10 | 0.527 | 76.35 | 7.76 | 38.78 | 0 |
| MAC-11 | 0.673 | 80.39 | 20.76 | 13.21 | 0 |
| MAC-12 | 0.591 | 80.36 | 16.77 | 18.15 | 0 |
| MAC-13 | 0.307 |  | 12.9 | 21.93 | 0 |
| MAC-14 | 1.5 | 82.16 | 15.73 | 42.64 | 0 |
| MAC-15 | 0.448 | 85.18 | 24.27 | 41.58 | 0 |
| MAC-16 | 0.597 | 85.56 | 15.14 | 29.34 | 0 |
| MAC-17 | 0.58 | 89.08 | 19.99 | 48.65 | 0 |
| MAC-18 | 0.322 | 77.24 | 16.23 | 28.95 | 0 |
| MAC-19 | 0.329 | 87.35 | 19.08 | 27.99 | 0 |
| MAC-20 | 5.3 | 85.25 | 11.74 | 71.69 | 0 |
| MAC-21 | 0.222 | 91.6 | 43.85 | NA | NA |
| MAC-22 | 0.333 | NA | 1.35 | NA | NA |
| MAC-23 | 0.815 | 72.45 | 19.99 | NA | NA |
| MAC-24 | 0.29 | 53.39 | 18.19 | NA | NA |
| MAC-25 | 0.359 | 61.81 | 24.32 | NA | NA |
| MAC-26 | 0.839 | 79.94 | 24.54 | NA | NA |
| MAC-27 | 0.524 | 70.47 | 18.82 | NA | NA |
| PAC-1 | 3.6 | 27.62 | 13.14 | 68.99 | 0 |
| PAC-2 | 0.104 | 5.78 | 16.08 | 48.39 | 0 |
| PAC-3 | 0.663 | 60.29 | 34.19 | 80.51 | 0 |
| PAC-4 | 0.531 | 68.7 | 17.93 | 83.03 | 0 |
| PAC-5 | 0.121 | 66.67 | 40.05 | 52.12 | 0 |
| PAC-6 | 2.79 | NA | 10.63 | 20.21 | 0 |
| PAC-7 | 0.625 | NA | 17.72 | 59.05 | 0 |
| PAC-8 | 3.25 | NA | 12.19 | 18.97 | 0 |
| PAC-9 | 0.912 | 27.58 | 35.23 | 61.75 | 0 |
| PAC-10 | 1.003 | 59.85 | 23.85 | 51.18 | 0 |
| PAC-11 | 0.088 | 57.79 | 32.54 | 64.4 | 0 |
| PAC-12 | 0.31 | 75.64 | 22.79 | 45.25 | 0 |
| PAC-13 | 0.157 | 75.82 | 31.03 | 70.78 | 0 |
| PAC-14 | 11.422 | NA | 24.28 | 9.07 | 0 |
| PAC-15 | 1.87 | NA | 38.65 | 23.72 | 0 |
| PAC-16 | 0.665 | 75.19 | 25.54 | 16.65 | 0 |
| PAC-17 | 0.202 | 53.3 | 34.92 | 58.07 | 0 |
| PAC-18 | 0.239 | 68.7 | 31.79 | 72.01 | 0 |
| PAC-19 | 0.356 | NA | 32.98 | NA | NA |
| PAC-20 | 0.507 | 68.7 | 32.6 | 47.04 | 0 |
| PAC-21 | 0.381 | 77.03 | 23.73 | 47.88 | 0 |
| PAC-22 | 0.812 | NA | 34.89 | 36.1 | 0 |
| PAC-23 | 1.725 | 65.06 | 20.69 | 4.77 | 0 |
| PAC-24 | 0.144 | NA | 14.15 | NA | NA |
| PAC-25 | 0.166 | 71.15 | 34.27 | 45.32 | 0 |
| PAC-26 | 0.105 | NA | 24.19 | NA | NA |
| PAC-27 | 0.237 | 82.73 | 27.11 | 87.27 | 0 |
| PAC-28 | 0.172 | 71.5 | 20 | 53.03 | 0 |
| PAC-29 | 0.519 | 86.17 | NA | 69.93 | 0 |
| PAC-30 | 9.615 | 72.46 | 21.25 | 49.54 | 0 |
| PAC-31 | 1.3 | 80.36 | 27.34 | 21.54 | 0 |
| PAC-32 | 0.378 | NA | NA | 25.67 | 0 |
| PAC-33 | 0.431 | 85.09 | 30.37 | 21.12 | 0 |
| PAC-34 | 0.472 | 83.59 | 31.61 | 57.9 | 0 |
| PAC-35 | 4.234 | 72.32 | NA | 34.37 | 0 |
| PAC-36 | 1.231 | 54.96 | NA | NA | 0 |
| PAC-37 | 1.539 | 68.24 | NA | 76.66 | 0 |
| PAC-38 | 0.103 | 64.4 | NA | 71.5 | 0 |
| PAC-39 | 0.14 | 80.47 | 13.9 | NA | 0 |
| PAC-40 | 0.104 | 33.98 | 26.28 | 45.44 | 0 |
| PAC-41 | 0.093 | 22.57 | 11.02 | 79.27 | 0 |
| PAC-42 | 0.213 | 89.1 | 20.94 | 89.97 | 0 |
| PAC-43 | 2.008 | 74.71 | 11.34 | 57.27 | 0 |
| PAC-44 | 1.627 | 70.32 | 17.56 | 21.81 | 0 |
| PAC-45 | 4.205 | 54.52 | 7.14 | 26.36 | 0 |
| PAC-46 | 1.111 | NA | 10.37 | 37.48 | 0 |
| PAC-47 | 0.133 | 79.11 | NA | 47.65 | 0 |
| PAC-48 | 3.25 | NA | NA | 50.34 | 0 |
| PAC-49 | 0.56 | 38.07 | NA | 32.06 | 0 |
| PAC-50 | 0.56 | 72.39 | NA | 29.61 | 0 |
| PAC-51 | 3.708 | 71.08 | 29.94 | 40.67 | 0 |
| PAC-52 | 48.362 | 69.11 | 20.35 | 28.09 | 0 |
| PAC-53 | 2.002 | 75.13 | 41.79 | 63.24 | 0 |
| PAC-54 | 3.629 | 47.87 | 38.6 | 28.86 | 0 |
| PAC-55 | 2.312 | 32.69 | 27.17 | 74.19 | 0 |
| PAC-56 | 1.824 | 65.49 | 25.94 | 36.63 | 0 |
| PAC-57 | 22.344 | 53.8 | 21.17 | 23.17 | 0 |
| PAC-58 | 9.427 | 66.78 | 16.3 | 21.82 | 0 |
| PAC-59 | 6.7 | 76.04 | 15.53 | 11.82 | 0 |
| PAC-60 | 0.815 | 38.96 | 8.65 | 55.75 | 0 |
| PAC-61 | 9.673 | 35.9 | 26.77 | 35.9 | 0 |
| PAC-62 | 2.528 | 25.63 | 25.87 | 25.63 | 0 |
| PAC-63 | 1.051 | 14.77 | 19.55 | 14.77 | 0 |
| PAC-64 | 0.826 | 23.66 | 30.41 | 23.66 | 0 |
| PAC-65 | 4.247 | 57.7 | 27.79 | 57.7 | 0 |
| PAC-66 | 5.524 | 52.71 | 24.74 | 52.71 | 0 |
| PAC-67 | 1.812 | 29.08 | 11.23 | 29.08 | 0 |
| PAC-68 | 1.444 | 45.37 | 23.77 | 45.37 | 0 |
| PAC-69 | 4.305 | 63.38 | 28.68 | 36.82 | 0 |
| PAC-70 | 5.228 | 75.22 | 19.23 | 39.44 | 0 |
| PAC-71 | 0.161 | 56.27 | 32.56 | 46.97 | 0 |
| PAC-72 | 0.503 | 54.02 | 26.97 | 41.35 | 0 |
| PAC-73 | 0.677 | 70.57 | 29.83 | 35.28 | 0 |
| PAC-74 | 0.357 | 73.9 | 31.42 | 56.56 | 0 |
| PAC-75 | 14.7 | 62.25 | 22.87 | 72.15 | 0 |
| PAC-76 | 8.9 | 76.03 | 25.9 | 42.27 | 0 |
| PAC-77 | 0.279 | 81.95 | 51.62 | 65.93 | 0 |
| PAC-78 | 0.556 | 85.98 | 42.08 | 74.7 | 0 |
| PAC-79 | 0.99 | 75.44 | 34.98 | 51.48 | 0 |
| PAC-80 | 1.051 | 88.7 | 25.58 | 59.66 | 0 |
| PAC-81 | 0.206 | 95.58 | 44.1 | 87.4 | 0 |
| PAC-82 | 16.625 | 73.47 | 12.54 | 37.37 | 0 |
| PAC-83 | 3.4 | 85.39 | 17.19 | 63.72 | 0 |
| PAC-84 | 2.502 | 48.29 | 16.54 | 60.32 | 0 |
| PAC-85 | 2.97 | 16.69 | 15.66 | 26.4 | 0 |
| PAC-86 | 5.648 | 58.81 | 8.92 | 51.07 | 0 |
| PAC-87 | 4.409 | NA | 16.45 | 39.72 | 0 |
| PAC-88 | 2.429 | 58.85 | 14.59 | 45.49 | 0 |
| PAC-89 | 4.126 | 27.9 | 12.83 | 29.42 | 0 |
| PAC-90 | 0.738 | 69.78 | 14.4 | 27.03 | 0 |
| PAC-91 | 11.1 | 58.38 | 13.15 | 45.21 | 0 |
| PAC-92 | 2 | 71.5 | NA | 73.55 | 0 |
| PAC-93 | 0.0158 | 57.47 | NA | 78.72 | 0 |
| PAC-94 | 0.178 | 52.86 | NA | 66.31 | 0 |
| PAC-95 | 5.4 | 64.09 | 9.68 | 67.2 | 0 |
| PAC-96 | 8.1 | 36.02 | 11.39 | 59.97 | 0 |
| PAC-97 | 1.4 | 70.03 | 16.29 | 44.95 | 0 |
| PAC-98 | 6.4 | 53.66 | 14.52 | 45.01 | 0 |
| ASC-1 | 9.7 | 52.96 | 24.32 | 17.01 | 0.98 |
| ASC-2 | 1.7 | 62.78 | 28.83 | 57.32 | 1.32 |
| ASC-3 | 5.6 | 65.98 | 41.84 | 11.79 | 12.05 |
| ASC-4 | 20.6 | 42.13 | 26.16 | 23.39 | 0.86 |
| ASC-5 | 0.3007889 | 26.15 | 47.25 | 52.251 | 7.6 |
| ASC-6 | 20.7 | 52.9 | 38.31 | 13.56 | 4.63 |
| ASC-7 | 8 | 60.35 | 28.58 | 51.78 | 4.23 |
| ASC-8 | 26.1 | 56.64 | 32.35 | 50.9 | 1.7 |
| ASC-9 | 22 | 65.51 | 27.89 | 34.08 | 7.46 |
| ASC-10 | 17.8 | 69.79 | 23.49 | 38.17 | 11.46 |
| ASC-11 | 0.9655288 | 69.01 | 46.86 | 71.46 | 0.09 |
| ASC-12 | 21.2 | NA | 41.4 | 9.54 | 5.35 |
| ASC-13 | 7.5 | 28.41 | 41.89 | 56.18 | 1.13 |
| ASC-14 | 15.5 | 50.07 | 29.67 | 28.61 | 0.35 |
| ASC-15 | 22 | 64.79 | NA | 19.67 | NA |
| ASC-16 | 22.8 | 69.09 | NA | 7.54 | NA |
| ASC-17 | 5.8 | 65.83 | 42.18 | 2.22 | 4.02 |
| ASC-18 | 35.8 | 70.81 | 34.95 | 43.99 | 0.04 |
| ASC-19 | 5.1 | 64.93 | 50 | 15.65 | 3.43 |
| ASC-20 | 4.8 | 68.54 | 45.47 | 68.3 | 4.8 |
| ASC-21 | 8.7 | 63.59 | 32.78 | 3.96 | 11.84 |
| ASC-22 | 96 | 70.3 | 35.05 | 17.89 | 21.13 |
| ASC-23 | 5.8 | 48.64 | 7.16 | 22.24 | NA |
| ASC-24 | 2.3 | 60.53 | 28.23 | NA | NA |
| ASC-25 | 1.3 | 47.04 | 15 | 33.85 | 5.8 |
| ASC-26 | 27.7 | 50.79 | 7.61 | 0.20126 | 14.68 |
| ASC-27 | 36.7 | 32.75 | 8.71 | 29.95 | 15.97 |

| Animal | Survival (days) | Metastasis |
| --- | --- | --- |
| KLS<sup>wt</sup>-SPC-1 | 142 |  |
| KLS<sup>wt</sup>-SPC-2 | 105 | Overt |
| KLS<sup>wt</sup>-SPC-3 | 123 |  |
| KLS<sup>wt</sup>-SPC-4 | 105 |  |
| KLS<sup>wt</sup>-SPC-5 | 120 |  |
| KLS<sup>wt</sup>-SPC-6 | 88 |  |
| KLS<sup>wt</sup>-SPC-7 | 122 | Overt |
| KLS<sup>wt</sup>-SPC-8 | 125 | Overt |
| KLS<sup>het</sup>-SPC-1 | 150 |  |
| KLS<sup>het</sup>-SPC-2 | 135 |  |
| KLS<sup>het</sup>-SPC-3 | 177 | Overt |
| KLS<sup>het</sup>-SPC-4 | 161 |  |
| KLS<sup>het</sup>-SPC-5 | 143 | Minimal |
| KLS<sup>het</sup>-SPC-6 | 114 | Overt |
| KLS<sup>het</sup>-SPC-7 | 133 | Overt |
| KLS<sup>het</sup>-SPC-8 | 90 | Overt |
| KLS<sup>het</sup>-SPC-9 | 160 |  |
| KLS<sup>het</sup>-SPC-10 | 129 | Minimal |
| KLS<sup>het</sup>-SPC-11 | 223 | Overt |
| KLS<sup>het</sup>-SPC-12 | 89 | Overt |
| KLS<sup>het</sup>-SPC-13 | 128 |  |
| KLS<sup>het</sup>-SPC-14 | 166 | Minimal |
| KLS<sup>het</sup>-SPC-15 | 131 | Minimal |
| KLS<sup>het</sup>-SPC-16 | 124 | Minimal |
| KLS<sup>het</sup>-SPC-17 | 104 | Minimal |
| KLS<sup>null</sup>-SPC-1 | 130 |  |
| KLS<sup>null</sup>-SPC-2 | 125 |  |
| KLS<sup>null</sup>-SPC-3 | 103 |  |
| KLS<sup>null</sup>-SPC-4 | 131 |  |
| KLS<sup>null</sup>-SPC-5 | 159 |  |
| KLS<sup>null</sup>-SPC-6 | 163 |  |
| KLS<sup>null</sup>-SPC-7 | 118 |  |
| KLS<sup>null</sup>-SPC-8 | 134 |  |
| KLS<sup>null</sup>-SPC-9 | 138 |  |
| KLS<sup>null</sup>-SPC-10 | 133 |  |
| KLS<sup>null</sup>-SPC-11 | 148 | Overt |
| KLS<sup>null</sup>-SPC-12 | 122 | Overt |
| KLS<sup>null</sup>-SPC-13 | 118 |  |
| KLS<sup>null</sup>-SPC-14 | 184 | Minimal |
| KLS<sup>null</sup>-SPC-15 | 132 |  |
| KLS<sup>null</sup>-SPC-16 | 142 |  |
| KLS<sup>null</sup>-SPC-17 | 87 | Minimal |
| KLS<sup>null</sup>-SPC-18 | 199 |  |
| KLS<sup>null</sup>-SPC-19 | 111 |  |
| KLS<sup>wt</sup>-CC10-1 | 96 |  |
| KLS<sup>wt</sup>-CC10-2 | 103 |  |
| KLS<sup>wt</sup>-CC10-3 | 76 |  |
| KLS<sup>wt</sup>-CC10-4 | 80 |  |
| KLS<sup>wt</sup>-CC10-5 | 113 |  |
| KLS<sup>wt</sup>-CC10-6 | 86 |  |
| KLS<sup>het</sup>-CC10-1 | 92 |  |
| KLS<sup>het</sup>-CC10-2 | 75 |  |
| KLS<sup>het</sup>-CC10-3 | 179 | SCC-like |
| KLS<sup>het</sup>-CC10-4 | 171 | AC |
| KLS<sup>het</sup>-CC10-5 | 115 |  |
| KLS<sup>het</sup>-CC10-6 | 97 |  |
| KLS<sup>het</sup>-CC10-7 | 66 |  |
| KLS<sup>het</sup>-CC10-8 | 114 |  |
| KLS<sup>het</sup>-CC10-9 | 103 | AC |
| KLS<sup>het</sup>-CC10-10 | 195 |  |
| KLS<sup>het</sup>-CC10-11 | 97 | SCC-like |
| KLS<sup>null</sup>-CC10-1 | 128 |  |
| KLS<sup>null</sup>-CC10-2 | 114 |  |
| KLS<sup>null</sup>-CC10-3 | 133 |  |
| KLS<sup>null</sup>-CC10-4 | 126 |  |
| KLS<sup>null</sup>-CC10-5 | 102 |  |
| KLS<sup>null</sup>-CC10-6 | 95 | SCC-like |
| KLS<sup>null</sup>-CC10-7 | 115 |  |
| KLS<sup>null</sup>-CC10-8 | 95 |  |
| KLS<sup>null</sup>-CC10-9 | 163 |  |
| KLS<sup>null</sup>-CC10-10 | 136 |  |
| KLS<sup>null</sup>-CC10-11 | 87 |  |
| KLS<sup>null</sup>-CC10-12 | 103 | SCC-like |
| KLS<sup>null</sup>-CC10-13 | 103 |  |

| **Animal** | **%IAC** | **%MAC** | **%PAC** |
| --- | --- | --- | --- |
| KLS<sup>wt</sup>-SPC-1 | 11.56 |  | 30.57 |
| KLS<sup>wt</sup>-SPC-2 | 0.1 |  | 28.52 |
| KLS<sup>wt</sup>-SPC-3 | 2.82 |  | 21.9 |
| KLS<sup>wt</sup>-SPC-4 | 5.02 |  | 31.39 |
| KLS<sup>wt</sup>-SPC-5 | 1.47 |  | 23.18 |
| KLS<sup>wt</sup>-SPC-6 | 0.68 |  | 29.36 |
| KLS<sup>wt</sup>-SPC-7 | 2.48 |  | 30.08 |
| KLS<sup>wt</sup>-SPC-8 | 0.22 |  | 34.59 |
| KLS<sup>het</sup>-SPC-1 | 3.56 |  | 22.53 |
| KLS<sup>het</sup>-SPC-2 | 5.5 |  | 9.73 |
| KLS<sup>het</sup>-SPC-3 | 4.88 |  | 26.89 |
| KLS<sup>het</sup>-SPC-4 | 4.19 |  | 15.08 |
| KLS<sup>het</sup>-SPC-5 | 8.85 |  | 12.07 |
| KLS<sup>het</sup>-SPC-6 | 6.91 |  | 9.46 |
| KLS<sup>het</sup>-SPC-7 | 11.58 |  | 5.27 |
| KLS<sup>het</sup>-SPC-8 | 11.75 |  | 25.601 |
| KLS<sup>het</sup>-SPC-9 | 15.56 |  | 20.5 |
| KLS<sup>het</sup>-SPC-10 | 6.28 |  | 2.32 |
| KLS<sup>het</sup>-SPC-11 | 8.85 |  | 6.7 |
| KLS<sup>het</sup>-SPC-12 | 3.69 |  | 11.56 |
| KLS<sup>het</sup>-SPC-13 | 12.38 |  | 19.11 |
| KLS<sup>het</sup>-SPC-14 | 17.9 |  | 0.41 |
| KLS<sup>null</sup>-SPC-1 | 4.58 |  | 20.57 |
| KLS<sup>null</sup>-SPC-2 | 15.76 |  | 2.22 |
| KLS<sup>null</sup>-SPC-3 | 8.91 |  | 1.67 |
| KLS<sup>null</sup>-SPC-4 | 10.22 |  | 2.93 |
| KLS<sup>null</sup>-SPC-5 | 9.91 |  | 0 |
| KLS<sup>null</sup>-SPC-6 | 8.31 |  | 27.81 |
| KLS<sup>null</sup>-SPC-7 | 17.79 |  | 1.17 |
| KLS<sup>null</sup>-SPC-8 | 27.413 |  | 8.36 |
| KLS<sup>null</sup>-SPC-9 | 14.78 |  | 18.12 |
| KLS<sup>null</sup>-SPC-10 | 15.92 |  | 0 |
| KLS<sup>null</sup>-SPC-11 | 22.57 |  | 36.49 |
| KLS<sup>null</sup>-SPC-12 | 16.2 |  | 6.52 |
| KLS<sup>null</sup>-SPC-13 | 24.95 |  | 3.89 |
| KLS<sup>wt</sup>-CC10-1 |  | 1.75 |  |
| KLS<sup>wt</sup>-CC10-2 |  | 2.99 |  |
| KLS<sup>wt</sup>-CC10-3 |  | 2.55 |  |
| KLS<sup>wt</sup>-CC10-4 |  | 0.25 |  |
| KLS<sup>wt</sup>-CC10-5 | 0.5 | 0.07 | 3.59 |
| KLS<sup>wt</sup>-CC10-6 |  | 3.18 |  |
| KLS<sup>het</sup>-CC10-1 |  | 0.16 |  |
| KLS<sup>het</sup>-CC10-2 | 0.11 | 0.33 |  |
| KLS<sup>het</sup>-CC10-3 | 1.59 | 0.86 | 2.99 |
| KLS<sup>het</sup>-CC10-4 | 0.43 | 12.51 | 4.99 |
| KLS<sup>het</sup>-CC10-5 | 0.81 | 2.36 | 4.22 |
| KLS<sup>het</sup>-CC10-6 | 0.15 | 1.17 |  |
| KLS<sup>het</sup>-CC10-7 | 4.59 | 24.09 |  |
| KLS<sup>het</sup>-CC10-8 | 0.04 | 1.35 |  |
| KLS<sup>het</sup>-CC10-9 | 0.1 | 0.063 |  |
| KLS<sup>het</sup>-CC10-10 | 3.24 |  | 7.3 |
| KLS<sup>het</sup>-CC10-11 |  |  |  |
| KLS<sup>null</sup>-CC10-1 |  |  |  |
| KLS<sup>null</sup>-CC10-2 | 0.08 | 0.7 |  |
| KLS<sup>null</sup>-CC10-3 | 0.21 | 2.75 |  |
| KLS<sup>null</sup>-CC10-4 | 1.42 |  |  |
| KLS<sup>null</sup>-CC10-5 | 3.87 | 0.03 |  |
| KLS<sup>null</sup>-CC10-6 |  | 0.13 | 1.27 |
| KLS<sup>null</sup>-CC10-7 | 0.19 | 0.83 |  |
| KLS<sup>null</sup>-CC10-8 | 0.13 |  |  |
| KLS<sup>null</sup>-CC10-9 | 1.44 | 0.19 | 1.07 |
| KLS<sup>null</sup>-CC10-10 | 0.19 | 0.62 |  |
| KLS<sup>null</sup>-CC10-11 | 0.59 |  |  |
| KLS<sup>null</sup>-CC10-12 | 0.13 | 0.04 |  |

| Animal | %ASC-papillary | %ASC-mucinous | SCC-like |
| --- | --- | --- | --- |
| KLS<sup>wt</sup>-CC10-1 | 15.35 | 22.33 |  |
| KLS<sup>wt</sup>-CC10-2 | 59.73 | 0 |  |
| KLS<sup>wt</sup>-CC10-3 | 56.01 | 7.44 |  |
| KLS<sup>wt</sup>-CC10-4 | 36.68 | 0 |  |
| KLS<sup>wt</sup>-CC10-5 | 37.14 | 0 |  |
| KLS<sup>wt</sup>-CC10-6 | 13.75 | 27.76 |  |
| KLS<sup>het</sup>-CC10-1 | 65.53 | 0 |  |
| KLS<sup>het</sup>-CC10-2 |  | 0 | 51.13 |
| KLS<sup>het</sup>-CC10-3 |  | 5.36 | 24.68 |
| KLS<sup>het</sup>-CC10-4 |  | 14.45 |  |
| KLS<sup>het</sup>-CC10-5 |  |  | 10.1 |
| KLS<sup>het</sup>-CC10-6 |  |  | 56.1 |
| KLS<sup>het</sup>-CC10-7 |  |  | 44.82 |
| KLS<sup>het</sup>-CC10-8 |  | 33.86 | 8.14 |
| KLS<sup>het</sup>-CC10-9 |  |  | 69.9 |
| KLS<sup>het</sup>-CC10-10 | 21.7 |  | 9.22 |
| KLS<sup>het</sup>-CC10-11 |  |  | 75.41 |
| KLS<sup>null</sup>-CC10-1 | 37.48 |  | 31.03 |
| KLS<sup>null</sup>-CC10-2 |  | 21.1 | 21.61 |
| KLS<sup>null</sup>-CC10-3 |  | 6.26 | 11.45 |
| KLS<sup>null</sup>-CC10-4 |  | 9.83 |  |
| KLS<sup>null</sup>-CC10-5 |  | 12.11 | 10.77 |
| KLS<sup>null</sup>-CC10-6 |  | 8.36 | 9.71 |
| KLS<sup>null</sup>-CC10-7 |  | 7.51 | 20.73 |
| KLS<sup>null</sup>-CC10-8 |  |  | 57.24 |
| KLS<sup>null</sup>-CC10-9 |  | 4.39 | 15.28 |
| KLS<sup>null</sup>-CC10-10 |  |  | 17.97 |
| KLS<sup>null</sup>-CC10-11 |  | 33.36 | 6.9 |
| KLS<sup>null</sup>-CC10-12 |  | 18.23 | 22.82 |

